# Unveiling the Molecular Mechanisms of the Type-IX Secretion System’s Response Regulator: Structural and Functional Insights

**DOI:** 10.1101/2024.05.15.594396

**Authors:** Anshu Saran, Hey-Min Kim, Ireland Manning, Mark A. Hancock, Claus Schmitz, Mariusz Madej, Jan Potempa, Maria Sola, Jean-François Trempe, Yongtao Zhu, Mary Ellen Davey, Natalie Zeytuni

**Affiliations:** Department of Anatomy and Cell Biology, McGill University, Montreal, Quebec, Canada. Address: 3640 Rue University, Montreal, Quebec, Canada H3A 0C7; Centre de Recherche en Biologie Structurale (CRBS), Montreal, Quebec, Canada; Department of Microbiology, The Forsyth Institute, Cambridge, MA 02142 USA; Department of Biological Sciences, Minnesota State University Mankato, Mankato, Minnesota, USA; Department of Pharmacology & Therapeutics, McGill University, Montreal, Quebec, Canada, H3G 1Y6; Department of Structural Biology, Molecular Biology Institute of Barcelona, Spanish Research Council, Barcelona Science Park, Barcelona, E-08028, Spain; Department of Microbiology, Faculty of Biochemistry, Biophysics and Biotechnology, Jagiellonian University, Kraków PL-30-387, Poland; Department of Oral Immunology and Infectious Diseases, School of Dentistry, University of Louisville, Louisville, Kentucky 40202, USA; Department of Biological Sciences, Xi’an Jiaotong-Liverpool University, Suzhou, Jiangsu, China

**Keywords:** Type-IX secretion system, response regulator, alkaline phosphatase, *Porphyromonas gingivalis*, bacterial pathogenicity

## Abstract

The Type-IX secretion system (T9SS) is a nanomachinery utilized by bacterial pathogens to facilitate infection. The system is regulated by a signaling cascade serving as its activation switch. A pivotal member in this cascade, the response regulator protein PorX, represents a promising drug target to prevent the secretion of virulence factors. Here, we provide a comprehensive characterization of PorX both *in vitro* and *in vivo*. First, our structural studies revealed PorX harbours a unique enzymatic effector domain, which, surprisingly, shares structural similarities with the alkaline phosphatase superfamily, involved in nucleotide and lipid signaling pathways. Importantly, such pathways have not been associated with the T9SS until now. Enzymatic characterization of PorX’s effector domain revealed a zinc-dependent phosphodiesterase activity, with active site dimensions suitable to accommodate a large substrate. Unlike typical response regulators that dimerize via their receiver domain upon phosphorylation, we found that zinc can also induce conformational changes and promote PorX’s dimerization via an unexpected interface. These findings suggest that PorX can serve as a cellular zinc sensor, broadening our understanding of its regulatory mechanisms. Despite the strict conservation of PorX in T9SS-utilizing bacteria, we demonstrate that PorX is essential for virulence factors secretion in *Porphyromonas gingivalis* and affects metabolic enzymes secretion in the non-pathogenic *Flavobacterium johnsoniae*, but not for the secretion of gliding adhesins. Overall, this study advances our structural and functional understanding of PorX, highlighting its potential as a druggable target for intervention strategies aimed at disrupting the T9SS and mitigating virulence in pathogenic species.

## 1. Introduction

*Porphyromonas gingivalis* is the keystone bacterial pathogen in periodontitis, a chronic inflammatory disease of the tooth supporting tissues (1). Periodontitis affects nearly half of the world’s population and ranks as the 6^th^ most prevalent disease globally. Disease manifestation results from bacterial plaque products directly affecting periodontium along with the induction of the host’s inflammatory and immune responses (2). Chronic exposure to these destructive processes leads to connective tissue destruction, periodontal pocket formation, alveolar bone loss, and even edentulation (3). In addition to *P. gingivalis’s* significant role in periodontitis, other epidemiological and experimental studies have linked its infection and virulence factors secretion to other systemic conditions, including cardiovascular diseases, rheumatoid arthritis, preterm low birth weight, non-alcoholic fatty liver disease, cancer, and Alzheimer’s disease (4–9).

*P. gingivalis* manifests its infection by the secretion of potent virulence factors that promote tissue invasion and destruction, as well as interference with the host’s defense systems (10). Among these virulence factors, a group of cysteine proteases, called gingipains, are considered essential to its survival and pathogenicity. These proteases with trypsin-like activity allow this bacterium to escape immune defenses via the cleavage and degradation of host receptors, immunomodulatory proteins, signalling pathway regulatory proteins and adhesion molecules (11). Gingipain secretion is mediated by a unique type-IX secretion system (T9SS) which includes two distinct steps. Initially, the gingipains are translocated across the bacterial inner membrane by the general Sec machinery. Then, once in the periplasm, the gingipains are directed by a conserved C-terminal domain to the specialized T9SS-translocon located at the outer membrane. Following the translocation across the outer membrane, the proteins are further processed to remove a conserved C-terminal shuttling domain that is simultaneously replaced with an anionic lipopolysaccharide and remains attached to the bacterial surface or are subsequently released into the extracellular milieu via outer membrane vesicles (12).

Aside from its role in periodontal and systemic disease manifestation in humans, the T9SS is the export apparatus for various virulence factors of different bacterial pathogens, favoring disease dissemination and manifestation in fish and birds (13–16). Notably, the T9SS also presents additional physiological roles in non-pathogenic bacterial species of the Fibrobacteres-Chlorobi-Bacteroidetes superphylum, as highlighted by Mark and Zhu (17). For instance, in the soil bacterium, *Flavobacterium johnsoniae*, the T9SS enables a unique mode of gliding motility through the secretion of specialized adhesins (18) and facilitates the degradation of polysaccharides such as chitin by specialized digestive enzymes secretion (19, 20).

To date, 18 conserved proteins were identified to govern the T9SS function (21). Deletion of any of these genes resulted in the accumulation of cargo proteins in the periplasm; a hallmark of a dysfunctional T9SS. Microbial and biochemical studies have further assigned these essential T9SS members into four functional groups involved in: (i) regulation, (ii) assembly of the core T9SS structures across the inner and outer membranes, (iii) cargo translocation across the outer membrane, and (iv) post-translational modification of the cargo protein (10). Among the regulatory group of protein components, conserved proteins consisting of a Two-Component System (TCS) and an additional sigma factor were identified. TCSs are the primary multistep signaling pathways in bacteria, and their minimal composition includes an input sensor histidine kinase (SHK) and an output response regulator (RR). Signal transduction via the TCS relies on phosphoryl-transfer reactions, and includes SHK autophosphorylation, phosphoryl group transfer to the RR, followed by RR dephosphorylation (22). The RR component comprises a conserved receiver domain that undergoes phosphorylation and variable output effector domain(s). While most of RR’s effector domains function as DNA-binding transcription factors, some exhibit RNA-binding, protein-binding, or even enzymatic activities (23).

Recent studies in *P. gingivalis* suggest that the perception of an unidentified signal in the periplasm leads to the autophosphorylation of the T9SS-SHK, PorY protein (24). The phosphorylated PorY then continues to trans-phosphorylate the T9SS-RR, PorX protein (24). Unlike canonical bacterial TCSs, PorY and PorX are not located within the same operon and are therefore considered orphan SHK and RR, respectively (22, 25–27). Notably, PorX also lacks the classical DNA-binding effector domain and instead possesses an enzymatic effector domain of an unknown function *in vivo*. To promote signal transduction through the pathway and regulate gene transcription, PorX interacts with the SigP transcription factor, which then binds directly to the promoter regions of T9SS genes (27, 28). This regulatory pathway model was further supported by the observation that either *porX* or *sigP* genes deletion resulted in the downregulation of T9SS components and activity, impaired processing of gingipains as well as decreased toxicity (29). Remarkably, this regulatory cascade not only directs the transcription levels of conserved T9SS machinery components but also directly modulates the inner membrane T9SS-rotary motor, driving virulence factors secretion across the outer membrane. This reported link between the regulatory pathway and the cargo secretion motor (part of core T9SS structures across the inner and outer membranes) relies solely on interaction between the PorX protein and the cytoplasmic domain of the rotary core component protein, PorL (30). It has been speculated that the PorX-PorL mode of interaction shares similarities with the regulation of the flagella rotary motor by the response regulator, CheY, in other bacteria (30).

While studies to date have highlighted PorX as the hub protein governing both transcription and motor functions of the entire T9SS, our recent investigation of PorX from *P. gingivalis* (PorX_PG_) unveiled its ability to hydrolyze crucial signaling molecules like cyclic and linear oligoadenylates (31). Despite these advancements, the precise role and mechanism by which this atypical response regulator mediates signal transduction through the cascade remains unknown. Here, we report the structure determination and functional analysis of the PorX protein from the gliding bacterium *F. johnsoniae* (PorX_FJ_) as a model system. Our crystal structures reveal that the PorX_FJ_ adopts a two-domain dimeric fold with a classical RR receiver domain’s fold and a unique Alkaline Phosphatase Superfamily (APS) domain connected by a helical linker region. Our functional characterization *in vitro* shows that PorX_FJ_ can be activated by dimerizing either upon phosphorylation at the receiver domain or by zinc binding at the APS domain, suggesting a plausible dual activation cellular signal. Next, we demonstrate a functional complementation between the PorX_FJ_ from the non-pathogenic *F. johnsoniae* bacterium to PorX_PG_ from the pathogenic *P. gingivalis*. Accordingly, we further translate the obtained structural and functional knowledge to investigate the PorX’s role in *P. gingivalis*. Accordingly, we find that conformational alterations in the APS domain induced by zinc, rather than its phosphodiesterase catalytic function, have a more pronounced impact on T9SS functionality in *P. gingivalis*. Together, our results mark a significant milestone in the characterization and understanding of the role of this critical hub protein in virulence factors secretion.

## 2. Methods

### Recombinant protein expression and purification in *Escherichia coli*

#### Construction of expression plasmids

The full-length Fjoh_2906 (UniProt A5FFU4) gene was amplified by polymerase chain reaction (PCR) from *Flavobacterium johnsoniae* ATCC 17061 strain UW101 genomic DNA. The restriction-free method (32) was used to insert the amplified gene into a modified pET28a(+) vector (Novagen). In the expression vector, the gene was fused in frame to an N-terminal 10x-His tag followed by a thrombin proteolytic site. Site-directed mutagenesis was performed by Gibson assembly (33). All primers and plasmids used in this study are listed in Table S1 and S2, respectively.

#### Bacterial cultivation and protein expression

*E. coli* BL21 strain cells harboring pET28a(+)-PorX_FJ_ variants (wildtype and mutants) were cultivated in an autoinduction medium (34) containing kanamycin (50 μg/ml) at 37°C for 8 h. The cultivation temperature was lowered to 22°C and expression continued for another 16 h. The cells were harvested by centrifugation at 6000 × *g* for 15 min at 4°C.

#### Recombinant protein purification

PorX_FJ_ variants were purified as previously described (35). Briefly, cell pellets were resuspended in buffer A (50 mM Tris pH 8, 300 mM NaCl and 10 mM imidazole) and incubated with DNase I (10 mg/ml) and protease inhibitor cocktail (Calbiochem) at 4°C. The cells were then disrupted by two cycles in a French press pressure cell at 172 MPa. Cell debris was removed by high-speed centrifugation at 270,000 × *g* for 1 h at 4°C. The supernatant was applied to a gravity Ni-NTA column (Bio-Rad Econo-Column chromatography column, Thermo Scientific HisPur Ni-NTA resin) pre-equilibrated with buffer A. The bound protein was washed with buffers B (20 mM Tris pH 8, 300 mM NaCl and 20 mM imidazole) and C (20 mM Tris pH 8, 1 M NaCl and 40 mM imidazole) and eluted with buffer D (20 mM Tris pH 8, 200 mM NaCl and 500 mM imidazole). To cleave the His-tag, bovine thrombin (Prolytix) was added to the eluted protein and the mixture was then dialyzed against buffer E (20 mM Tris pH 8, 200 mM NaCl) for 16 h at 4°C. The protein was then applied onto a size exclusion column (Superdex 200 16/60 GL, Cytivia) pre-equilibrated with buffer E. The purified proteins were concentrated to ∼25 mg/ml and flash-frozen in liquid nitrogen.

### Crystallization and structure determination

Wild-type PorX_FJ_ and the T271V mutant were each crystallized using the sitting drop vapour diffusion method at 25°C. In brief, 0.2 μl of 5 mg/ml protein solution was mixed with an equal volume of precipitant solution. Native PorX_FJ_ was crystallized in two different conditions. Detailed crystalization conditions and individual data collection parameters for each crystal are listed in Supplementary Table S3. For phasing, the PorX_FJ_ primitive orthorhombic crystal form was soaked for 15 sec in a precipitant solution containing 0.5 M sodium bromide and multiwavelength anomalous diffraction (MAD) datasets were collected. All datasets were reduced and scaled using the HKL2000 suite (36) and phases were obtained using CRANK2 (37). The initial model was manually built in Coot (38) according to the electron density map calculated from data collected at the peak wavelength of bromide. This initial atomic model was used as a template for molecular replacement in Phaser (39) against all other datasets. The final models were manually edited in Coot (38) and refined by Refmac5 (40). For R_free_ calculations, 5% of the data were excluded. All structural figures were prepared using Chimera (41). Data collection and refinement parameters are listed in Supplementary Table S4. Structures have been submitted to the Protein Data Bank (accession codes 8TEF, 8TED, 8TFF, 8TFM, 8THP).

### Isothermal titration calorimetry (ITC)

Binding affinity measurements were performed at 20°C using an isothermal titration calorimeter (Microcal iTC200, Malvern). 50 μM PorX_FJ_ was diluted in buffer E and placed in the sample cell. Buffer E supplemented with 750 μM zinc chloride was placed in the syringe and titrated into the protein samples. Each titration was 10 μl in volume and lasted for 10.3 sec followed by an equilibration period of 240 s.

### Inductively coupled plasma mass spectrometry (ICP-MS)

PorX_FJ_ variants and method blanks, were digested in PTFE vessels using trace metal grade concentrated nitric acid at room temperature overnight, followed by 2 h at 90°C. Digestates were diluted with deionized water to 2 % w/w nitric acid and the concentrations of Mg, Mn, Co, Zn, and Cd were determined by ICP-MS. A Thermo X-Series 2 ICP-MS with collision cell technology (CCT) and chilled spray chamber was used in kinetic energy discrimination (KED) mode with 8 % hydrogen in helium to reduce interferences. The protein-divalent metal cation stoichiometries were determined by dividing the total number of the protein and each divalent metal cation number of moles.

### Phosphodiesterase activity assay *in vitro*

PorX_FJ_ variants (2.5 μM wild-type or mutants) in buffer G (50 mM Tris pH 8, 150 mM sodium chloride) were screened for their phosphodiesterase activity. Similarly, the phosphorylated variants were incubated in buffer G supplemented with 20 mM AcP and 5 mM MgCl_2_ for 1 h at 37°C prior to measurements. For metal screening, wild-type PorX_FJ_ was incubated in buffer G alone or supplemented with 0.5 mM ZnCl_2_, CuCl_2_, MnCl_2_, MgCl_2,_ or CaCl_2_. For activity pH screening, phosphorylated and non-phosphorylated PorX_FJ_ were incubated in 150 mM NaCl, 3 μM ZnCl_2_, supplemented with 50 mM acetic acid (pH 6), 50 mM Tris (pH 7-9) or CAPS buffer (10–11) for 30 min at 37°C. The catalysis of bis(4-nitrophenyl) phosphate (bis-*p*NPP), *p*-nitrophenyl phosphate (*p*NPP) or *p*-nitrophenyl sulphate (*p*NPS) (0-10 mM) and the formation of *p*-nitrophenol product were monitored at 405 nm on a Synergy H1 microplate reader (BioTek). The bis-*p*NPP reaction was monitored for 2 h at 37°C, while the *p*NPP and *p*NPS reactions were monitored for 3 days at 37°C. All assays were performed in quadruplicates. However, the absorbance measurements of the T271V mutant could not be obtained for bis-*p*NPP concentrations exceeding 5 mM.

### Protein phosphorylation assay *in vitro*

For intact protein mass spectrometry, purified PorX_FJ_ variants were incubated at 100 μM in buffer F (25 mM Tris pH 7.5, 10 mM MgCl_2_, 1 mM MnCl_2_, 2 mM dithiothreitol, 5 mM AcP, 0.01 mM sodium orthovanadate). After 20 min at 37°C (or 1h at room temperature for PorX_FJ_ D54A/T271V due to precipitation issues), the reactions were quenched by flash-freezing in liquid nitrogen.

### Intact protein liquid chromatography mass spectrometry (LC-MS)

As previously described (42–44), protein samples were diluted to 0.1 mg/mL in 0.1% (v/v) formic acid before 1 µg was injected on a Dionex Ultimate 3000 UHPLC system at 40 µl/min using a Waters Acquity BEH C4 BEH column (300Å, 1.7 µM, 1 × 100 mm). The resulting eluate (5 min wash with 4% (v/v) acetonitrile followed by 15 min gradient of 4–90% (v/v) acetonitrile in 0.1% (v/v) formic acid) was analyzed on an Impact II QTOF mass spectrometer (Bruker Daltonics) equipped with an Apollo II ion funnel electrospray ionization source and Bruker otofControl v4.0 / DataAnalysis v4.3 software. Data were acquired in positive-ion profile mode using a capillary voltage of 4,500 V and dry nitrogen heated at 200°C. Total ion chromatograms were used to determine where the protein eluted and spectra were summed over the entire elution peak. Multiple charged ion species were deconvoluted at 5,000 resolution using the maximum entropy method.

### Protein dimerization assay *in vitro*

PorX_FJ_ variants (100 μM wildtype or mutants) were incubated in buffer E supplemented with either 300 μM ZnCl_2_ or 20 mM acetyl phosphate (AcP) and 5 mM MgCl_2_ for 1 h at room temperature. The mixture was then subjected to size exclusion chromatography (Superdex 200 increase 10/300 GL or Superdex 75 increase 10/300 GL), pre-equilibrated with buffer E.

### Functional characterization in *P. gingivalis*

#### Bacterial strains, plasmids, and growth conditions

The wild-type strain used in this study was *P. gingivalis* W50. The wild-type bacterium and mutants were propagated from −80°C freezer stocks and grown anaerobically at 37°C for 3-5 days on agar plates containing Trypticase Soy Broth (TSB) (Becton, Dickinson and Company, Franklin Lakes, NJ, USA) supplemented with 5 µg/ml hemin, 1 µg/ml menadione, and 5% defibrinated sheep blood (BAPHK) (Northeast Laboratory Services, Winslow, ME, USA). The atmosphere of the anaerobic chamber contained a mixture of 5% hydrogen, 10% carbon dioxide and 85% nitrogen. The bacterial colonies were used for making starter cultures in TSB liquid media supplemented with 5 µg/ml hemin and 1 µg/ml menadione that were then grown anaerobically at 37°C without shaking.

#### Construction of the deletion mutants in *P. gingivalis*

Mutant strain W50 *ΔporX_PG_* was generated as previously described (45). In order to construct the *ΔporX_PG_* mutant, 1 kb long regions both upstream and downstream of *porX_PG_*, were amplified from W50 genomic DNA and the erythromycin resistance gene (*emrF*) was amplified from plasmid pVA2198 by PCR. The three amplicons were purified and combined using the NEBuilder HiFi DNA Assembly Master Mix (New England BioLabs, Ipswich, MA, USA) according to the instructions provided by the manufacturer. The final product was transformed into *P. gingivalis* by electroporation. The transformed cells containing the erythromycin resistance gene in the site of *porX_PG_* were selected by growth on BAPHK medium containing 10 μg/ml erythromycin.

To complement the *porX_PG_* deletion mutant, *porX_PG_* and *porX_FJ_* were cloned separately into plasmid pT-COW and placed under the control of a low-level constitutive promoter [groES (PG0521) promoter region], generating pT-groES-*porX_PG_* and pT-groES-*porX_FJ_* respectively. To generate all truncations and point mutations of *PorX_PG_*, the pT-groES-*porX_PG_* plasmid was used. The pT-groES-*porX_PG_* constructions with different mutations on the *porX_PG_* region were then transformed into *E. coli* S17-1 cells for the conjugation with *ΔporX_PG_* mutant. All complemented strains were generated by conjugation as previously described (46). In brief, BAPHK containing tetracycline (1 µg/ml) was used to select for pT-groES-*porX_PG_* containing *P. gingivalis* strains, and gentamicin (200 µg/mL) was used to counter-select the *E. coli* S17-1 donor. Transconjugants were obtained after 7 days of anaerobic incubation and the transconjugants were isolated, verified by PCR, and maintained on BAPHK containing tetracycline (1 µg/mL). All primers and plasmids used in this study are listed in Table S1 and S2, respectively.

#### Gingipain enzymatic activity assay

Wild-type *P. gingivalis* W50, PorX null mutant cells carrying empty pT-COW plasmids, and all pT-groES-*porX_PG_* constructs carrying the different mutations of *porX_PG_* were cultivated to OD_600_ of 1.0. The cultures were harvested and screened for their gingipain activities. Arginine-gingipain (Rgp) and Lysine-gingipain (Kgp) assays were carried out according to previously described methods (47) with slight modifications. For the Rgp assay, 10 μl of liquid culture was mixed with 170 μl of buffer H (200 mM Tris HCl, pH 7.6, 150 mM NaCl, 5 mM CaCl_2_, 0.02% NaN_3_ and 20 mM L-cysteine previously neutralized with 8 M NaOH in a 9:1 ratio). In the case of the Kgp assay, 50 μl liquid culture was mixed with 130 μl buffer H. The mixtures were incubated at 37°C for 10 min followed by the addition of 1mM substrate (Nα-Benzoyl-L-arginine 4-nitroanilide hydrochloride for the Rgp assay and 2-acetamido-6-amino-*N*-(4-nitrophenyl) hexanamide for the Kgp assay). The formation of p-nitroaniline product was measured at 405 nm at 1 min intervals for 30 min with constant shaking. All assays were performed in quadruplicates.

#### Expression levels validation by Western blot

Wild-type *P. gingivalis* W50 and plasmid complemented null *porX* strains were cultivated to OD_600_ of 1.0. The cultures were harvested and resuspended in phosphate buffer (PBS) supplemented with 2 mM of Nα-tosyl-l-lysine chloromethyl ketone hydrochloride (Sigma) and protease inhibitor cocktail (Calbiochem) to inactivate all gingipains prior to lysis. The resuspended cells were sonicated and boiled in SDS-PAGE sample buffer without dithiothreitol (DTT) for 3 min. The samples were boiled again for additional 3 min following the addtion of DTT to a final concntration of 20 mM. Samples were centrifuged briefly at 13,000 × g for 1 min to remove particulates and the supernatant separated on SDS-PAGE. Proteins were subsequently electro-transferred onto a nitrocellulose membranes and blocked in 5% [w:v] BSA/PBS solution overnight. Rabbit polyclonal anti-PorX_PG_ antibodies were manufacterd by Proteogenix (Schiltigheim, France) using recombinant PorX as an antigen. PorX_PG_ was detected using a rabbit of anti-PorX_PG_ (1 μg/ml) in PBST (PBS supplemented with 0.1% Tween 20) for 90 min. Membranes were washed three times with TTBS before being probed for 60 min with a 1:10,000 dilution of a polyclonal goat anti-rabbit horseradish peroxidase-conjugated secondary antibody (Invitrogen). Development was carried out using the ECL Western blot substrate kit according to the manufacturer’s instructions (Millipore). The biotinylated inner membrane-associated protein MmdC was used as a loading control (48) and detected by a 1:4,000 dilution of streptavidin conjugated to horseradish peroxidase (Thermo Scientific).

### Functional characterization in *F. johnsoniae*

#### Bacterial strains, plasmids, and growth conditions

*F. johnsoniae* UW101 (49, 50) was the wild-type strain used in this study. *F. johnsoniae* strains were grown at 25° in CYE liquid (51) or TYES (52). For solid media, 15 g agar was added per liter. *E. coli* strains were grown in Luria-Bertani medium (LB) at 37°C (53). For most experiments, *F. johnsoniae* strains were propagated from −80 °C glycerol stocks on CYE agar and incubated for 48-72 h at 25°C before they were used as starter cultures. All primers and plasmids used in this study are listed in Table S1 and S2, respectively. Antibiotics were used at the following concentrations when needed: ampicillin, 100 μg/ml; erythromycin, 100 μg/ml; kanamycin, 35 μg/ml; and tetracycline, 20 μg/ml unless indicated otherwise.

#### Construction of the deletion mutants in *F. johnsoniae*

For deletion of *porX_FJ_* (Fjoh_2906), a 2.0-kbp fragment spanning part of Fjoh_2905 and the first 105 bp of *porX_FJ_* was amplified using primers 0076 (introducing a BamHI site) and 0077 (introducing an XbaI site), and the Phusion DNA polymerase (Thermo Fisher Scientific, Waltham, MA). The fragment was digested with BamHI and XbaI and ligated into pYT313 (54), which had been digested with the same enzymes, to generate pIM03. A 1.9-kbp fragment spanning Fjoh_2907, Fjoh_2908, and the last 45 bp of *porX_FJ_* was amplified using primers 0167 (introducing an XbaI site) and 0079 (introducing a SalI site). The fragment was cloned into XbaI and SalI sites of pIM03 to generate the deletion construct pIM06. Plasmid pIM06 was introduced into *F. johnsoniae* UW101 by triparental conjugation as previously described (55). An erythromycin-resistant clone was streaked for isolation, and grown overnight in CYE liquid with shaking at 25°C in the absence of antibiotics. These cells were plated on CYE agar containing 5% sucrose and incubated at 25°C for 2-3 days. Sucrose-resistant colonies were streaked for isolation and screened by PCR using primers 0086 and 0087, which flank *porX_FJ_*, to identify the deletion mutant CJ4057. The same procedure was used to delete *gldKLMNO* using the plasmids and primers listed in Table S1 and Table S2, respectively.

#### Complementation of the *porX* deletion mutant

Primers 0086 (introducing a BamHI site) and 0087 (introducing an XbaI site) were used to amplify a 2009-bp fragment spanning *porX* with its putative promoter. The fragment was digested with BamHI and XbaI and ligated into pCP23 (56), which had been digested with the same enzymes, to generate pIM10. The plasmid was transferred to the *porX* mutant by triparental conjugation. Tetracycline was used for screening of the complemented colonies.

#### Analysis of cell motility

Cells of the wild type *F. johnsoniae* and *porX_FJ_* deletion mutant were grown for 17 h at 25°C in motility medium (57) without shaking. Tunnel slides were constructed using double stick tape, glass microscope slides, and glass coverslips, as previously described (58). Ten microliters of cultures were introduced into the tunnel slides, incubated for 5 min, and observed for motility using an Olympus CX41 phase-contrast microscope at room temperature (22°C). Images were recorded using a Moment CMOS camera and analyzed using Ocular (Teledyne Photometrics, Tuscon, AZ). Rainbow traces of cell movements were made using Fiji (https://imagej.net/) and the macro–Color FootPrint (59).

#### Growth of *F. johnsoniae* on chitin

Chitin powder (practical grade from shrimp shells; Sigma C7170) was prepared as a 1% slurry essentially as described previously (60) and used as a stock solution to prepare the chitin media. Cells of *F. johnsoniae* were streaked on fresh TYES agar and incubated at 25°C for 2 d. Cells were scraped off the plates, suspended in 1 ml Stanier medium (61), pelleted by centrifugation at 4,200 × g for 3 min to remove the residual TYES medium, and resuspended in Stanier medium to a concentration (OD600) of 1.0. Then 0.1 ml of the inoculation cell suspension was introduced into 50 ml of Stanier medium supplemented with 0.05% (w/v) chitin in 250-ml flasks, and incubated at 25°C with shaking (200 rpm). 2.5 μg/ml of tetracycline was added for growth of the complemented strain. At various times, 1 ml samples were removed. Cells and residual chitin were collected by centrifugation at 17,000 × g for 10 min. Growth was assessed by determining the total cellular protein in the pellets using the Bradford assay as previously described (62).

## 3. Results

PorX is a conserved protein found in some pathogenic and non-pathogenic bacterial species commonly known to utilize their T9SS for protein cargo secretion. In an effort to determine the structure of PorX, we have applied an ortholog screening approach (35). Ortholog screening is a well-established strategy to improve the expression, purification, and structural determination of challenging proteins (63). Among the different orthologs screened (35), we identified two distinctive conditions promoting the crystallization of the full-length PorX from *F. johnsoniae* (PorX_FJ_) in its non-phosphorylated apo-form. Heavy atom derivative (bromide) soaking, followed by a multi-wavelength anomalous diffraction data collection, were used to retrieve the missing phase information, and solve the high resolution structure of PorX_FJ_ (Table S3 and Table S4).

### Overall structure

Although the purified non-phosphorylated PorX_FJ_ was found to be monomeric in solution by size exclusion chromatography (35), both crystal forms presented an identical dimeric assembly within their asymmetric units (Fig. 1). The primitive orthorhombic crystal form gave rise to a single dimeric assembly (2 monomers in the asymmetric unit), while the primitive monoclinic crystal form gave rise to two identical dimeric assemblies (4 monomers in the asymmetric unit) (Fig. S1A-C). All PorX_FJ_ dimers adopt an intertwined assembly, where each monomer is symmetrically wrapped around the other monomer, forming an overall ‘X’-like shape (Fig. 1A). The formation of these identical assemblies (Cα’s RMSD = 0.98-2.35 Å^2^) suggests that the observed dimer represents a true functional state that was unlikely driven by crystal packing.

**Fig. 1.**
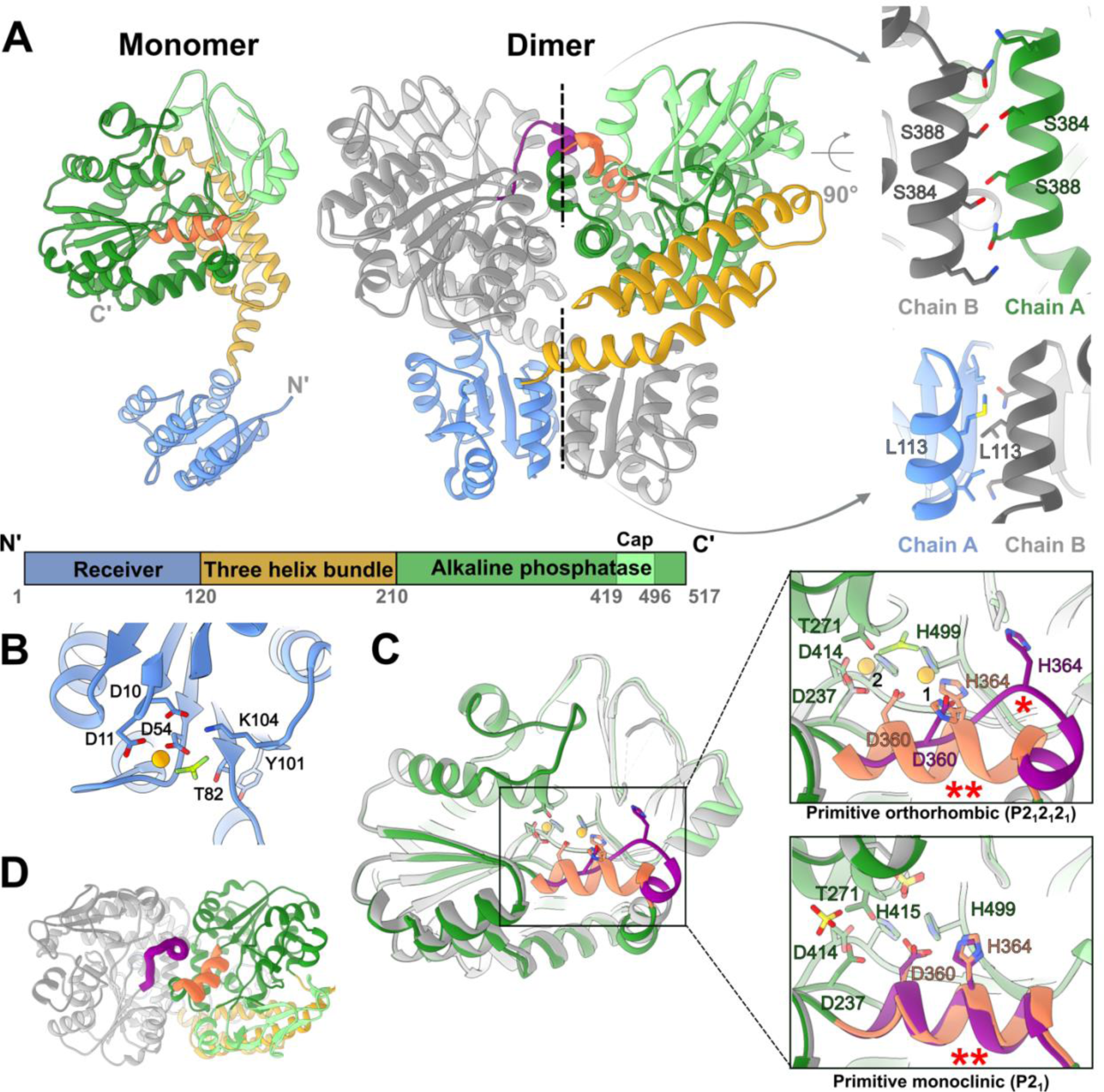
Crystal structures of PorX_FJ_. **(A)** Monomeric and dimeric assembly of PorX_FJ_. One monomer is coloured according to the different domains while the other monomer is in gray. The two dimeric interfaces are labelled by dashed lines and their zoom-in views highlight crucial residues at these dimerization interfaces. **(B)** Phosphorylation site at the REC domain. The conserved Asp54 coordinates both the phosphomimetic BeF_3_ molecule and a divalent cation presented as an orange sphere. Additional residues involved in coordination and stabilization of the phosphorylation site are presented. **(C)** Structural variation of the APS domain in the different monomers. Overlay of different APS domain conformers demonstrate the observed conformational change in residues 358-369. In the zoom-in views of the catalytic site, key residues associated with the coordination of two divalent cations (binding sites labelled as 1 and 2) and substrate binding/catalysis are presented. The substrate-mimicking BeF_3_ ion (shades of green) or sulfate ion (yellow), specifically identified in different PorX_FJ_ crystals is presented separately. The alternative conformation of residues 358-369, labelled by ^(^*^)^, disrupts the optimal positioning of the conserved divalent cation coordinating residues, Asp360 and His364. The helical conformation of residues 358-369, which maintains the ideal position of Asp360 and His364 is labelled by ^(^**^)^. **(D)** A top view of the APS domains in the PorX_FJ_ dimer demonstrate a conformational change in residues 358-369 (colored in purple and orange in each monomer), located in close proximity to the APS domain dimerization interface.

### Monomeric structure

Each PorX_FJ_ monomer adopts a two-lobed fold containing an N-terminal receiver (REC) domain (residues 1-120), followed by a three-helix bundle (THB) domain (residues 121-209) and a C-terminal alkaline phosphatase superfamily (APS) domain (residues 210-517) (Fig. 1A). All of the determined PorX_FJ_ monomers exhibit a high degree of structural similarity with a Cα RMSD ranging from 0.64 to 2.05 Å^2^. The minute structural variations observed primarily stem from the flexibility in the positioning of the REC domain relative to the THB and APS domains (Fig. S1D-E).

The highly conserved fold of the REC domain includes five ⍺-helices (⍺1-⍺5) that surround five central parallel β-strands (β1-β5). Within the REC domain, the surface exposed phosphorylation site involves residues Asp10, Asp11, Asp54, Thr82 and Lys104 that together coordinate the binding of a divalent cation and the phosphate moiety (Fig. 1B).

The THB domain interconnects the REC domain and the APS domain as it gives rise to the formation of the ‘X’-shaped dimer (Fig. 1A). In particular, the entire THB domain curves ∼75° away from the REC domain via its ⍺1 helix that packs against the REC domain of the adjacent monomer, while utilizing its ⍺2 and ⍺3 to pack against and support the subsequent APS domain.

At the C-terminal of PorX_FJ_, the APS domain comprises a β-sheet flanked by ⍺-helices (canonical (⍺/β)_6_ APS fold) and an additional β-strand rich capping region, also known as the cap sub-domain (Fig. 1A). The APS catalytic site is characterized by two divalent cations essential for catalysis and located within a shallow groove, where the substrate is predicted to bind. The two cations are coordinated by seven highly conserved residues: Asp360, His364 and His499 coordinates cation binding at the first site (Zn1), while Asp237, Asp414 and His415 and the catalytic nucleophile Thr271 coordinates cation binding at the second site (Zn2, sites nomenclature according to (67)) (Fig. 1D). Within the dimer, both APS catalytic sites are facing each other and are in close proximity to the dimeric interface (Fig. 1A). The divalent cation’s identity was confirmed by a co-crystallization in the presence of zinc and anomalous diffraction analysis (Table S3 and Table S4). In addition, zinc-binding was confirmed by inductively coupled plasma mass spectrometry (ICP-MS) revealing 2.02 zinc ions bound per each monomer. Zinc binding affinity was further analyzed by Isothermal Titration Calorimetry (ITC) and conformed to a two-site binding model with the following stoichiometries (N) and equilibrium dissociation constants (K_D_): N_1_= 1.09 ± 0.04 and K_D1_=180.83 ± 14.93 nM; N_2_=1.19 ± 0.04 and K_D2_= 59.17 ± 13.81 nM (Fig. S2).

### Dimeric interfaces

Two major interfaces mediate the formation of the intertwined dimer; the first is formed by the two identical REC domains (952 Å^2^), while the other is formed by the two APS domains (∼1000 Å^2^) (Fig. 1A). Notably, the phosphorylation and activation of canonical RRs lead to their dimerization via local conformational changes in two conserved REC domain residues. The induced dimerization then renders the downstream activities of the RR (23). Multiple RR crystal structures have unambiguously demonstrated that conserved threonine and tyrosine residues, also known as the TY-pair of the REC domain, undergo conformational changes upon the domain’s phosphorylation and are indicative of its activation state (23). In the non-phosphorylated state, the side chain of the conserved threonine faces the solvent, while in the phosphorylated state, it is re-oriented towards the phosphorylation site, thus creating a space to accommodate the conserved tyrosine side chain. These local conformational changes give rise to the formation of the REC domain dimers. Here, the positioning of the REC domain’s TY-pair (Thr82 and Tyr101) clearly indicates that the formed dimer represents the activated state (Fig. S3A).

To further assess if the observed dimer truly represents the phosphorylated/activated state, we have co-crystallized PorX_FJ_ with the phosphate analog, BeF_3_ (Table S3 and Table S4). In the presence of BeF_3_, PorX_FJ_ continued to adopt the same ‘X’-shaped dimer with a clear density that can be attributed to the analog observed at the REC domain’s phosphorylation site (Fig. 1B, Fig. S3B). In addition, in the primitive monoclinic crystal form, spherical densities, in dimensions suitable to accommodate sulfate ion (mimicking phosphate ion) originating from the crystallization condition, were clearly observed in the REC domain phosphorylation site (Fig. S3B). These observed phosphate-analog densities at the REC phosphorylation site were also found in similar positions to the phosphate moiety observed in the determined structure of the phosphorylated Spo0A response regulator (64). Taken together, we conclude that the observed dimeric fold truly represents the functional activated state.

The two APS domains give rise to the second dimerization interface of the intertwined dimer (Fig. 1A). The APS dimeric interface comprises symmetrical interaction between helix ⍺17 (residues 378-393) and residues 358-369 (Fig. 1A and Fig. S4). Despite the overall conserved fold of the APS domain (RMSD = 0.88 Å^2^), one area of significant difference specifically involves residues 358-369 (Fig. 1C-D). In the primitive orthorhombic crystal form, residues 358-369 adopt a helical conformation in one monomer while presenting an extended coil conformation and an additional helical turn at the N-terminal of helix ⍺16 in the second monomer (Fig. 1C-D). Moreover, in one of the monomers of the primitive monoclinic crystal form, residues 358-369 were flexible and their backbone could not be traced in the density map, while in the other monomers they adopted a helical conformation, further establishing the dynamic nature of this area of the protein (Fig. 1D). Global comparison of the APS dimerization interface demonstrates that the 358-369 helical-extended coil conformation dyad gives rise to a total of 1080 Å^2^ surface area while the helical-helical conformations dyad interface is only 993 Å^2^, suggesting the helical-extended loop conformation to be more stable.

Within each dimer, both APS active sites are located in close proximity to the dimeric interaction interface (Fig. 1A, C). Therefore, changes in the conformational state of the 358-369 region can significantly impact the zinc ion coordination of catalytic site (Fig. 1D). In particular, only the helical conformation of residues 358-369 can support the proper coordination of the Zn1 by the conserved Asp360 and His364 residues, whereas in the extended coil conformation, these residues are facing away from the APS catalytic site (Fig. 1D). Additional support to the link between metal coordination and residues 358-369 conformation is provided by the T271V APS catalytically inactive mutant structure (Table S3 and Table S4). In the T271V structure, a density corresponding to a divalent cation could only be observed in Zn1 site (coordinated by Asp360, His364 and His499) and not in the Zn2 site (coordinated by Thr271, Asp237, Asp414 and His415) (Fig. S5A), while residues 358-369 of both monomers adopted a helical conformation instead of the helical-extended coil dyad observed in the same primitive orthorhombic crystal form of the wildtype. Moreover, ICP-MS measurements revealed that the T271V mutant had a reduced ability to bind zinc with only 1.22 ions per monomer, instead of two seen by the wildtype. The ICP-MS measurements of the D360A/H364A double mutant further demonstrated a reduction in zinc binding capacity (0.63 zinc ions per monomer), suggesting that the zinc binding at the Zn1 site stabilizes the binding in the Zn2 site. Interestingly, abolishing the APS dimeric interface by the S348A/S388E double mutant also reduced the zinc-binding capacity to 1.03 zinc ions per monomer, thus, further establishing the link between zinc-binding at the APS domain and its dimer interface. Based on the observed conformational dynamics and the association with zinc binding, we speculate that this region might provide an additional level of regulation to the catalytic activity of the APS domain.

### Insights for substrate binding and catalytic activity of the APS domain

Structural comparison of the APS domain revealed a remarkable fold similarity to the alkaline phosphatase superfamily that can hydrolyze P-O, S-O, or P-C bonds while possessing a bi-metal zinc coordination architecture at their active site. Among these identified homologs, PhnA is a bacterial enzyme involved in a metabolic pathway that converts phosphonates to phosphorus and directly catalyzes phosphonoacetate to acetate and inorganic phosphate (65) (Fig. 2A). The other homologs belong to the EctoNucleotide Pyrophosphatase/Phosphodiesterase (ENPP) family (66–72) (Fig. 2A). The ENPP family members (ENPP1-7) can be found in both bacteria and eukaryotes, where they participate in different cellular processes, including nucleotide hydrolysis, lipid metabolism and associated signaling transduction pathways (73). Apart from the highly conserved residues associated with the catalytic mechanism and the two zinc ions coordination (Thr271, Asp360, His364, His499, Asp237, Asp414 and His415), the potential substrate binding pockets displayed some degree of variation depending on their specific substrate. In particular, ENPPs that are involved in oligonucleotide hydrolysis displayed the most extensive cavities while PhnA displayed the smallest cavity, compatible with the small size of the phosphonoacetate substrate (Fig. 2B). Here, our structural comparisons revealed that the APS site binding pocket of PorX_FJ_ is extensive and is, therefore, likely to bind a large substrate (Fig. 2B). In support of this notion, in the primitive monoclinic crystal form, two phospho-mimicking sulfate ions were identified and modeled at PorX_FJ_’s APS binding cavity (Fig. 2B, Fig. S5B). One sulfate ion was modelled next to the catalytic Thr271, while the other was located in an auxiliary binding cleft and stabilized by Arg275, Lys329 and Asn357. These findings further suggest that PorX_FJ_’s potential ligand might not only be large but also include poly-phosphate moieties such as cyclic di-nucleotides or polynucleotides. Notably, in the primitive orthorhombic crystal form, that was co-crystallized with the phospho-mimetic ion (Table S3 and Table S4), the corresponding density of a single BeF_3_ molecule was clearly observed in the catalytic site (in close proximity to the catalytic T271) but not in the auxiliary binding site (Fig. 2B, Fig. S5C). Notably, the spatial positions of the phosphate analogs identified in our PorX_FJ_ structures align closely with those observed in the recently determined crystallographic structure of catalytically inactive T272V variant of PorX_PG_ bound to phosphoguanylyl-(3′→5′)-guanosine (pGpG) (Fig. 2B, Fig. S6). However, our attempts to co-crystalize the catalytically inactive T271V variant of PorX_FJ_ with pGpG did not reveal the corresponding density in the APS catalytic site (Fig. S5A).

**Fig. 2.**
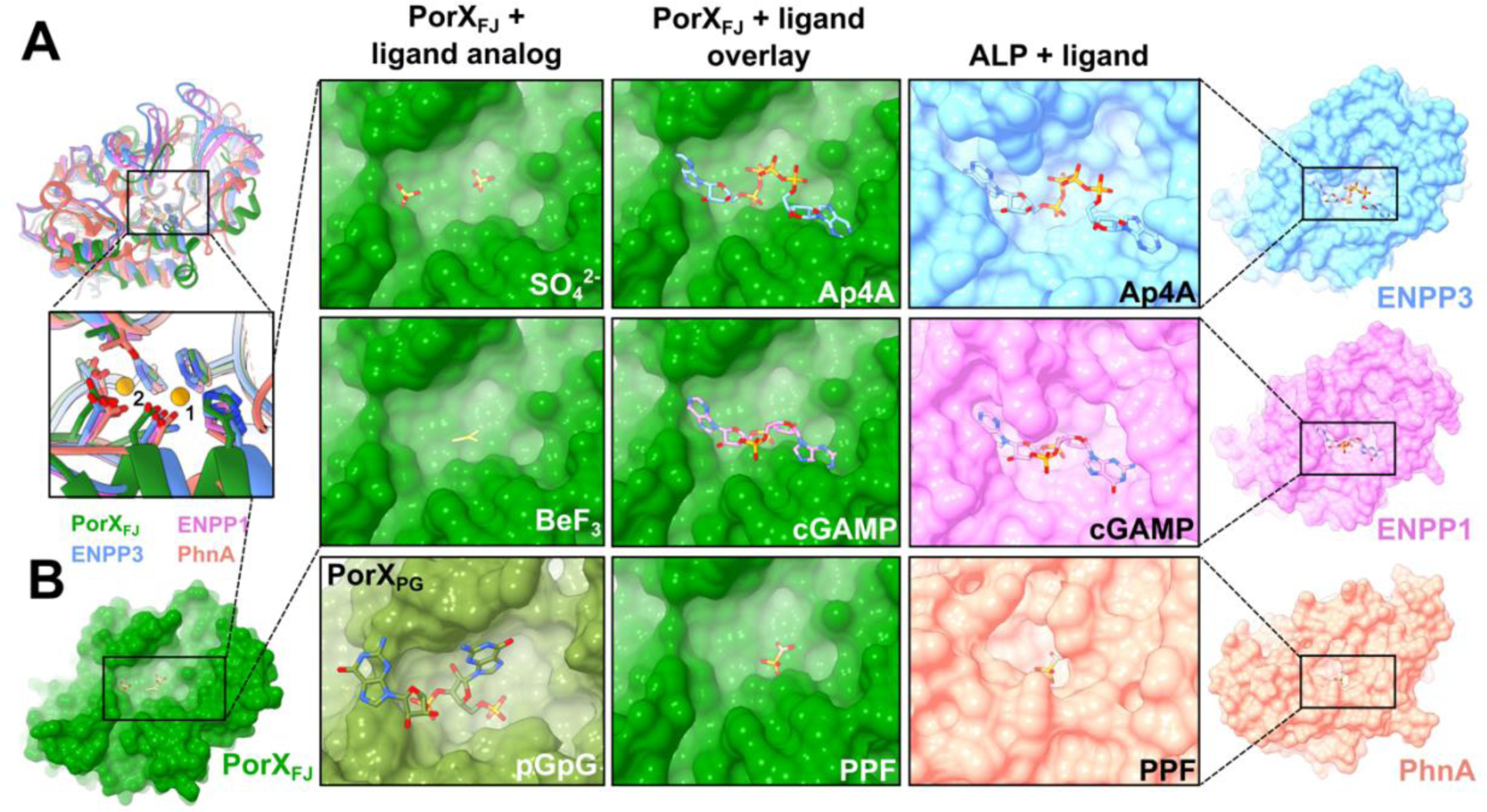
Structural similarities of PorX_FJ_‘s APS domain points towards its substrate specificity. **(A)** Overlay of the APS domains of PorX_FJ_, ENPP3, ENPP1 and PhnA demonstrates their conserved overall fold. Zoom-in view of their catalytic sites including the conserved bi-metal coordination, substrate binding and catalysis residues. **(B)** Surface representation of APS domains catalytic pockets exhibiting a clear correlation between the ligand and binding pocket dimensions. The APS domain of PorX_FJ_ with its determined ligand analogs (SO_4_^2-^ and BeF_3_) in green. PorX_PG_ with 5′-phosphoguanylyl-(3′→5′)-guanosine (pGpG) bound in olive green (PDB ID: 7PVK). The ENPP3 with Bis(adenosine)-5’-tetraphosphate (Ap4A) polynucleotide bound (PDB ID: 6F2Y) in blue, the ENPP1 with Adenosine-Guanosine-3’,3’-cyclic monophosphate (cGAMP) bound (PDB ID: 6AEL) in pink, while PhNA with phosphonoformate (PPF) bound (PDB ID: 1EI6) is in orange. Overlay of PorX_FJ_ with the ligand-bound ENPPs or PhnA suggests that the APS domain of PorX_FJ_ can accommodate a large poly-phosphate substrate.

Albeit our unsuccessful attempts to capture the structure of PorX_FJ_ in complex with cyclic or polynucleotides, we have confirmed its enzymatic activity by a phosphodiesterase assay. In the assay, full length PorX_FJ_ and the THB+APS truncation mutant (where the REC domain was truncated) were able to hydrolyze the phosphodiesterase substrate analog bis(*p*-nitrophenyl) phosphate (bis-*p*NPP) only in the presence of zinc and not by other divalent cations (Fig. 3A). The phosphodiesterase activity was pH dependent with an optimal pH in the range of 8-9, similar to the APS superfamily (Fig. 3B). Notably, significant reduction in hydrolysis rate was observed for (*p*-nitrophenyl) phosphate while no hydrolysis was observed for the (*p*-nitrophenyl) sulfate, providing hints towards substrate specificity (Fig. S7). Mutating the catalytic residues T271V, the metal coordinating residues D360A/H364A, and the APS dimerization interface residues S384A/S388E, abolished the observed phosphodiesterase activity against bis-*p*NPP (Fig. 3C). The phosphodiesterase assay further established the importance of zinc coordination and dimerization of the APS domain for catalytic activity.

**Fig. 3.**
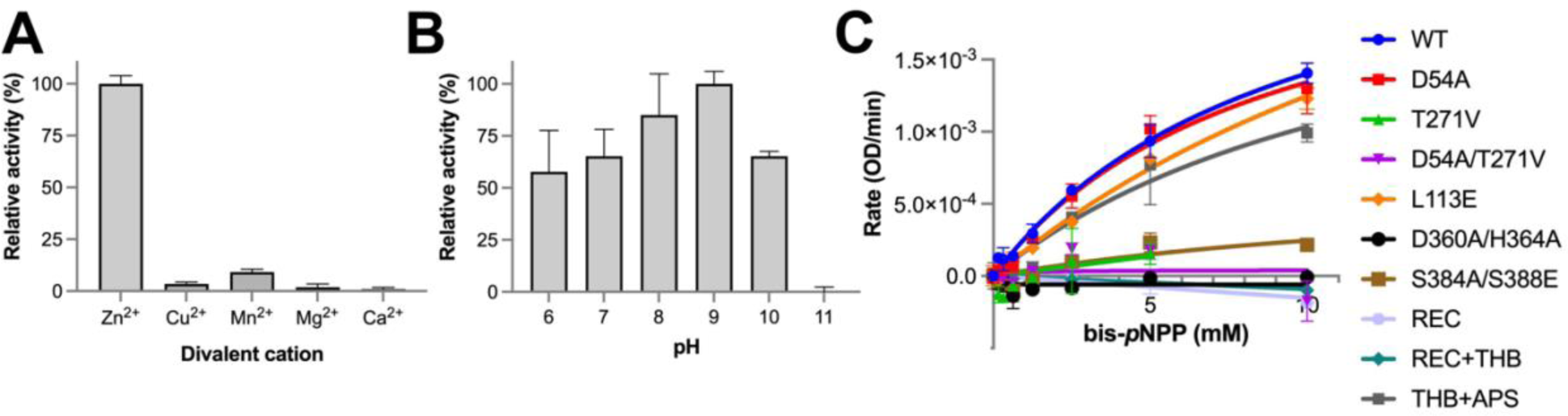
Phosphodiesterase activity of PorX_FJ_. **(A)** Divalent cation and **(B)** pH screening for the identification of optimum conditions of phosphodiesterase activity using bis-*p*NPP as a substrate. All percentage values in the pH and metal screens are normalized to pH 9 and Zn, respectively. **(C)** Phosphodiesterase kinetics of wild type PorX_FJ_ and its mutants at increasing concentrations of bis-*p*NPP.

### *In vitro* phosphorylation and subsequent dimerization of PorX_FJ_ variants

In TCS, canonical response regulators can be phosphorylated *in vitro* by incubation with the potent phosphodonor acetyl phosphate (AcP). Phosphorylation of the conserved Asp residue at the response regulator’s REC domain often leads to its dimerization and further induces its downstream function. Here, incubation with AcP in the presence of Mg^2+^ also led to PorX_FJ_ phosphorylation, as confirmed by intact protein mass spectrometry (LC-MS) experiments (Fig. 4, Table S5). Surprisingly, these LC-MS experiments revealed that apart from the predicted Asp54 phosphorylation site, AcP can also phosphorylate Thr271 in PorX_FJ_ (Fig. 4, Table S5). Using the REC domain’s phosphorylation-null mutant (D54A), the APS domain’s catalytically inactive mutant (T271V) and their double mutant (D54A/T271V), we confirmed that these residues can specifically undergo phosphorylation in PorX_FJ_ (Fig. 4, Table S5, Fig. S8). Interestingly, incubation of the wild-type PorX_FJ_ and certain mutants with AcP (apart from the D54A-phosphorylation null mutant), led to an 18 Da mass decrease in both the non-phosphorylated and phosphorylated populations (Fig. 4, Table S5). We hypothesized that the observed mass reduction resulted from a dehydration reaction linked to the formation of an internal peptide bond due to non-productive AcP phosphorylation (Fig. S9). Since the proposed cyclization reaction likely involved an amine group replacing one of the oxygens of Asp54, our molecular modeling identified a lysine residue (Lys104) in close proximity to Asp54 (Fig. 1B). Subsequent, intact LC-MS for the purified K104A mutant eliminated the dehydration reaction (Fig. 4, Table S5), thus validating our proposed alternative cyclization as an alternative AcP mechanism *in vitro*.

**Fig. 4.**
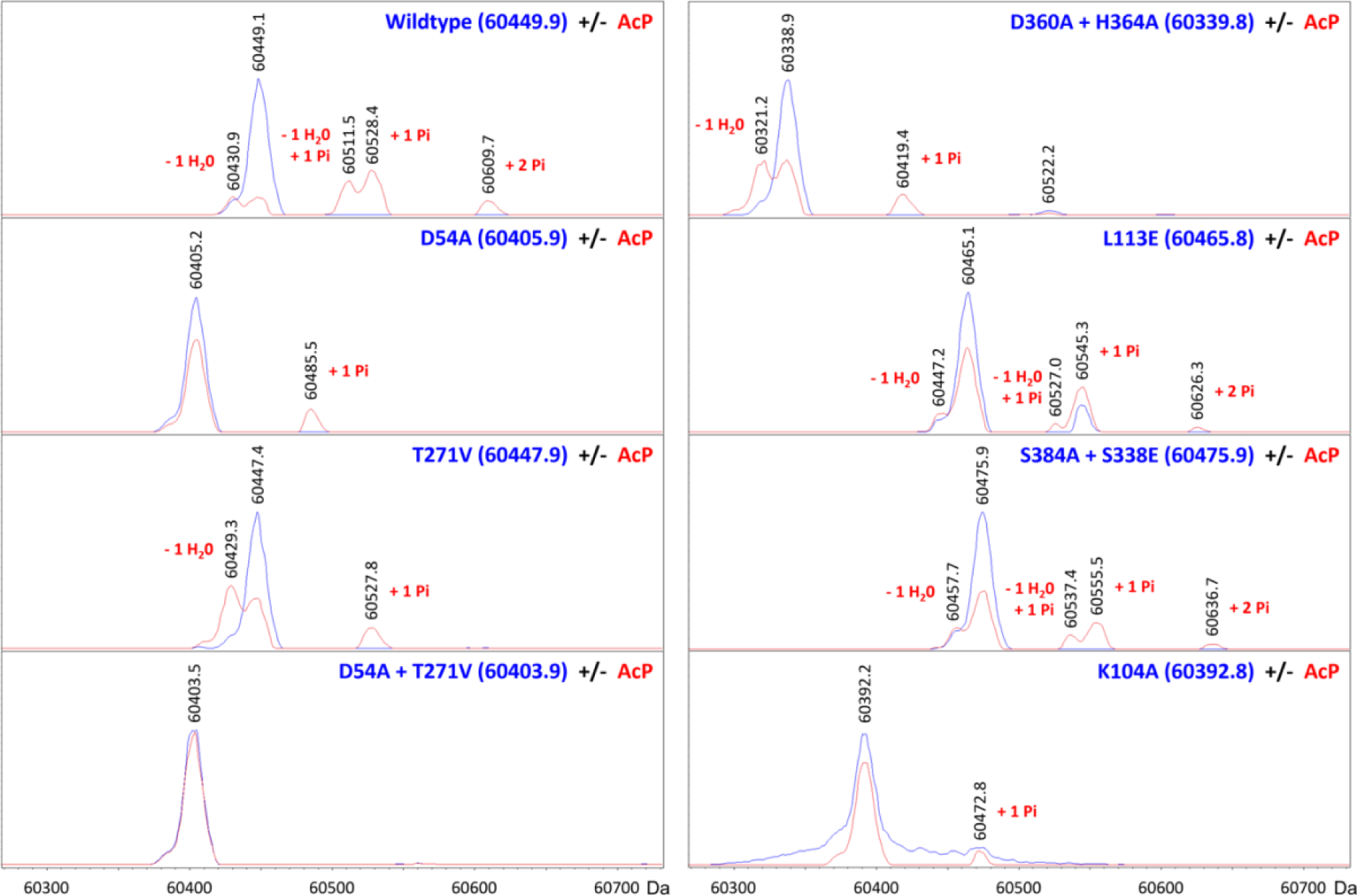
Intact protein LC-MS analyses of PorX_FJ_ point mutants in the absence or presence of phosphorylation *in vitro*. The blue and red spectra correspond to non-phosphorylated and phosphorylated versions of the protein, respectively. Comparing the theoretical and observed masses (inset blue and black numbers, respectively, in Da) showed that the acetyl phosphate (AcP) phosphodonor promotes the phosphorylation of Asp54 and/or Thr271 (“+1 Pi” or “+2 Pi” labels). Alternatively, AcP was found to induce cyclization of Asp54 and Lys104, resulting in a dehydration reaction (“-1 H_2_0” labels) and subsequent phosphorylation at Thr271 only (“-1 H_2_0, +1 Pi” labels).

Incubation with AcP in the presence of Mg^2+^ also led to PorX_FJ_ dimerization, as revealed by size exclusion chromatography (Fig. 5). Notably, incubation with Mg^2+^ alone did not result in protein dimerization. Mutation of the phosphorylation site in the REC domain (i.e., D54A) also eliminated the AcP-induced dimerization (Fig. 5). Interestingly, the L113E mutation, located at the REC domain dimerization interface, failed to dimerize in the presence of AcP, and thus, revealing the crucial cross-talk between phosphorylation and dimerization of the REC domain (Fig. 5).

**Fig. 5.**
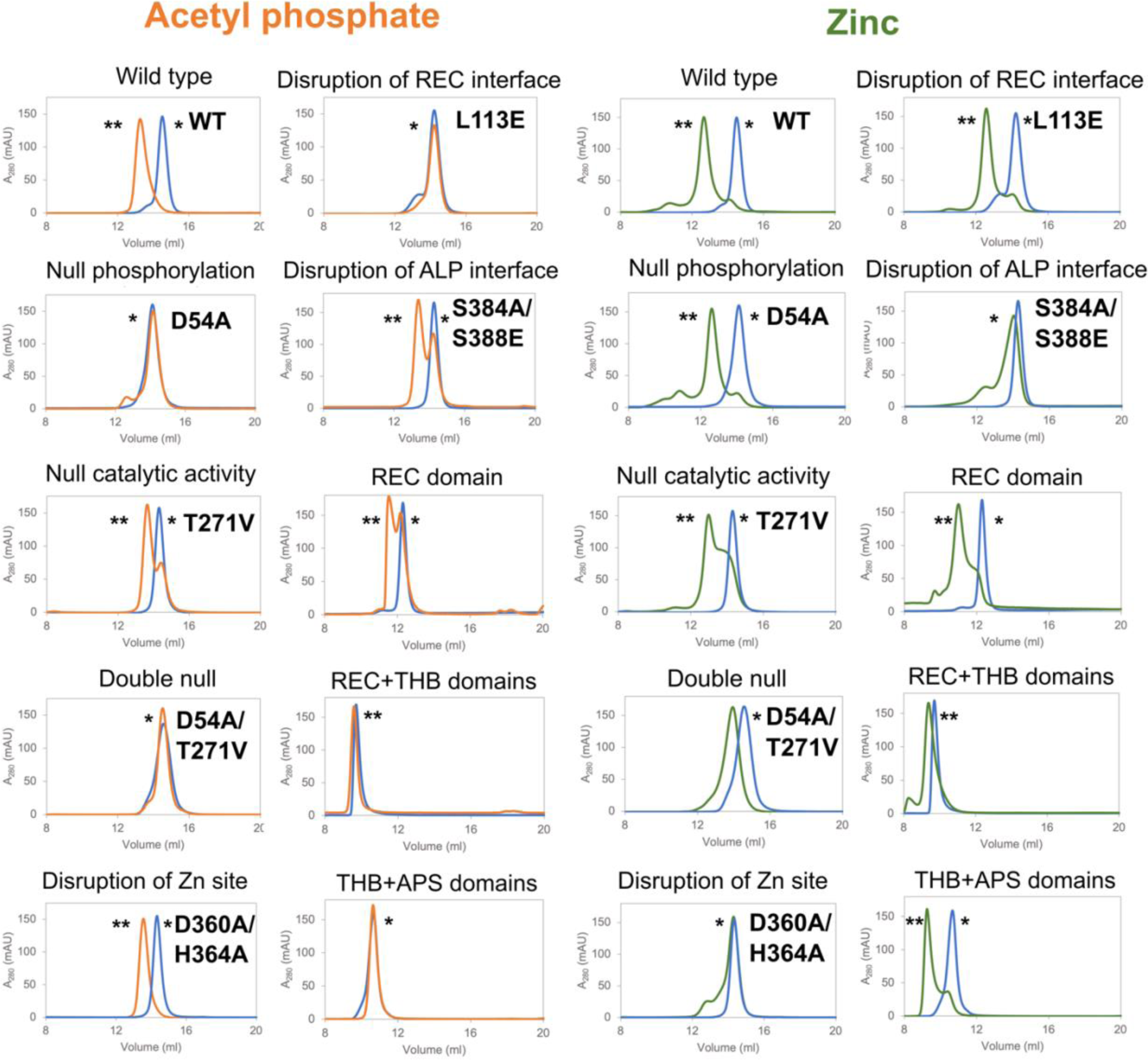
*In vitro* dimerization of PorX_FJ_ variants analysed by size exclusion chromatography. PorX_FJ_ proteins were incubated with acetyl phosphate in the presence of magnesium, or with zinc only. The blue, orange and green chromatograms refer to the proteins in the absence of any dimerizing agent, proteins with AcP or Zn, respectively. The peaks representing the monomeric and dimeric states of the protein are labelled by (*) and (**), respectively. WT and point mutation variants (∼61 kDa) were analysed by the Superdex200inc 10/300 GL column, while the truncation variants (REC domain: ∼14 kDa, REC+THB domains: ∼24 kDa and THB+APS domains: ∼46 kDa) were analysed by Superdex75inc 10/300 GL as per the column’s ability to better separate proteins with lower molecular weights.

### Zinc induced dimerization

In addition to zinc’s essential role for the catalytic activity of the APS domain, incubation of PorX_FJ_ with zinc resulted in a clear shift of its monomeric population to dimeric, even in the absence of AcP (Fig. 5). Notably, zinc-induced dimerization is independent of the REC domain ability to phosphorylate, as the D54A null phosphorylation, the L113E dimerization interface and the REC domain only truncation variants were still able to dimerize in the presence of zinc (Fig. 5). In contrast, mutations in the APS domain’s catalytic site and dimerization interface led to significant alterations in the zinc-induced dimerization ability (Fig. 5), which stands in good agreement with their reduced ability to bind zinc as demonstrated by the ICP-MS measurements. For example, the catalytically inactive T271V variant exhibited a mixed population of both monomers and dimers while the zinc coordination D360A/H364A double mutant remained mostly monomeric upon incubation with zinc (Fig. 5). Abolishing the APS dimerization interface in the S384A/S388E double mutant also resulted in a significant shift towards the monomeric population (Fig. 5). Interestingly, both the D54A and the T271V individual phosphorylation-null and catalytically inactive variants sustained a restricted ability to dimerize in the presence of zinc (Fig. 5). However, the double catalytic mutant D54A/T271V was not able to dimerize properly (Fig. 5), therefore, suggesting a cumulative effect of these mutations in comparison to their single catalytic mutants.

### Functional complementation and analysis of PorX in the pathogenic *P. gingivalis*

Deletion of the *porX* gene in *P. gingivalis* leads to a dysfunctional T9SS and subsequently severely impacts gingipains secretion (24, 28). Therefore, monitoring the gingipains secretion by measuring their amidolytic activity serves as a sensitive probe for the T9SS function in *P. gingivalis* (10, 74, 75). Since the orthologous proteins PorX_FJ_ and PorX_PG_ share 51% sequence identity and 71% sequence similarity, we hypothesized that PorX_FJ_ could functionally complement the Δ*porX* deletion strain of *P. gingivalis* (Fig. S4). Using the gingipains amidolytic activity assay (47), we demonstrated that the plasmid complementation of PorX_FJ_ can functionally substitute PorX_PG_, in the background of the Δ*porX_PG_* deletion strain (Fig. 6A). The observed functional complementation of PorX_FJ_ in *P. gingivalis* provided crucial evidence to our scientific approach as it validates the use of insights obtained from the PorX_FJ_ and the recently determined PorX_PG_ structures (31) to characterize the function of PorX_PG_ in the pathogenic *P. gingivalis.* As such, domains and key residues revealed by our reported PorX_FJ_ high-resolution structure determination are likely to share a similar function and be translatable for PorX_PG_, where their effect on virulence factors secretion can be further assessed *in vivo*.

**Fig. 6.**
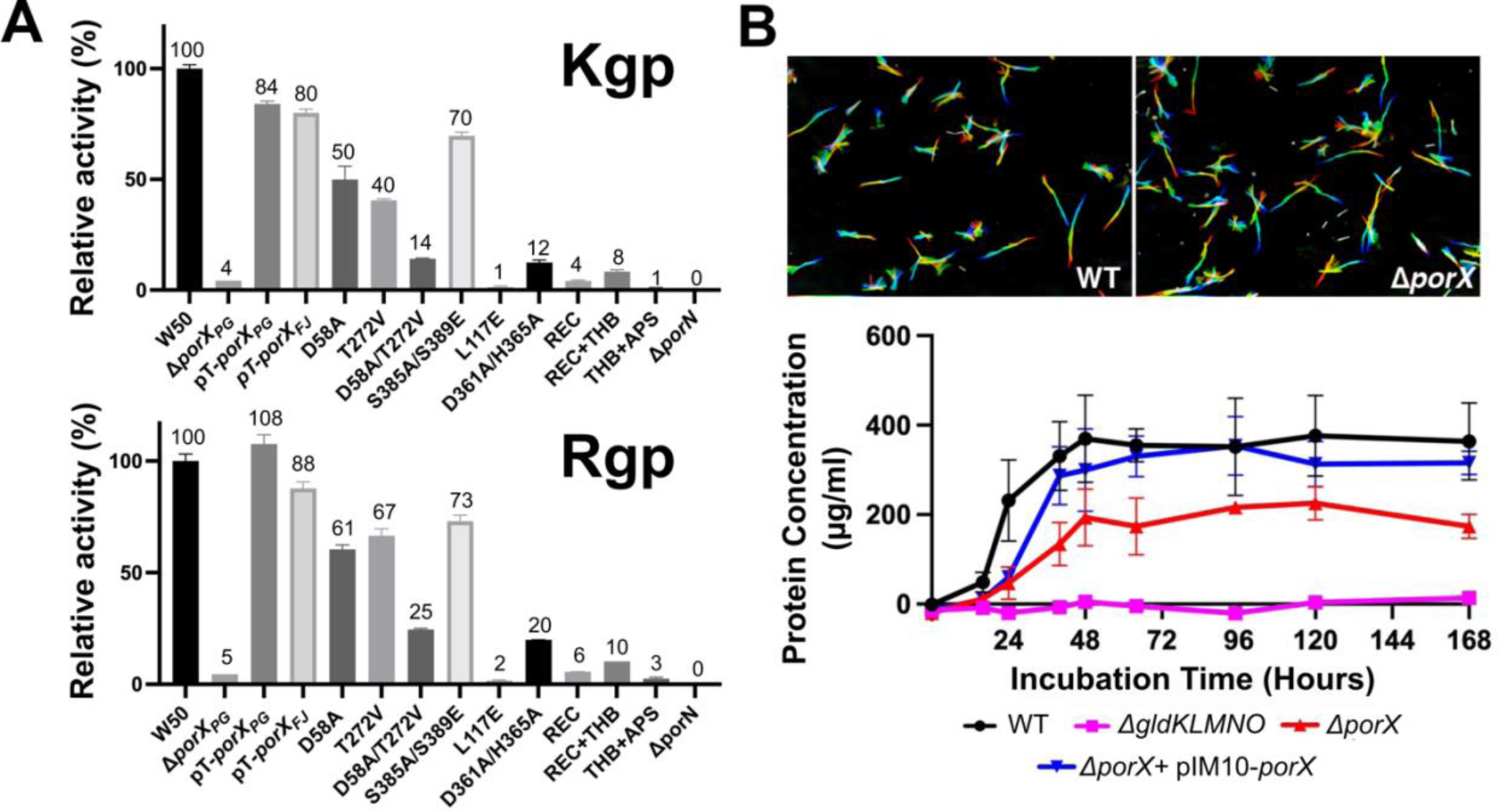
*In vivo* characterization of PorX in *P. gingivalis* and *F. johnsoniae*. **(A)** Gingipains enzymatic activity assay of *P. gingivalis* strain W50 containing different mutants of PorX. W50 and Δ*porX_PG_* refers to wild-type and PorX null mutant cells respectively, both carrying empty pT-COW plasmids. The pT-*porX_PG_* and pT-*porX_FJ_* represents complementation of Δ*porX*_PG_ pT-COW cells with wild-type PorX_PG_ and PorX_FJ_ respectively. All other point mutations and truncations include the complementation of Δ*porX_PG_* pT-COW cells with PorX_PG_ carrying the relevant mutation. Experiments were performed in triplicate and error bars indicate standard deviations. **(B)** *Top -* Gliding motility assay for the wild-type and Δ*porX* null mutant in *F. johnsoniae*. A series of 180 frames were collected and colored from red (time zero) to yellow, green, cyan, and finally blue and then integrated into one image, resulting in rainbow traces of gliding cells. Scale bar at lower right indicates 20 µm and applies to all panels. *Bottom* - Growth curve of the wild-type and Δ*porX* null mutant *F. johnsoniae* on a media containing chitin as the sole carbon source. Growth was presented as μg cellular protein/ml. Growth experiments were performed in triplicate and error bars indicate standard deviations. The T9SS null mutant, Δ*gldKLMNO,* that is unable to grow on chitin was used as the negative control. pIM10 contains the wild type *porXFJ* and was used to complement the Δ*porX* mutant.

First, we investigated the role of the PorX’s receiver and effector domains on virulence factors secretion in *P. gingivalis*. Here, our gingipains amidolytic activity assay revealed that the deletion of either the APS domain (PorX_PG_ Δresidues 208-517), the THB and the APS domains (PorX_PG_ Δresidues 126-517), or the REC domain (PorX_PG_ Δresidues 1-129), negatively affected the gingipain activity and thus, indicating a reduced gingipains secretion by the T9SS (Fig. 6A). The reduced amidolytic activity observed suggests that the standalone domains cannot restore the entire T9SS function and associated virulence factors secretion in *P. gingivalis* and that all domains must be present to support downstream function (Fig. 6A). Next, we assessed the effect of key residues mutations, initially identified in PorX_FJ,_ converted into their corresponding residues in PorX_PG_. We utilized Western blotting analyses (by anti-PorX_PG_ antibody) to confirm that the expression levels of all these point mutation variants were similar to the wild type PorX_PG_ and the effects seen are truly due to functional alteration (Fig. S10). Surprisingly, the gingipains amidolytic activities of the single phosphorylation-null and catalytically inactive variants (REC domain: D54A_FJ_ to D58A_PG_, or APS domain: T271V_FJ_ to T272V_PG_) were only partially reduced when compared to the wild type (Kgp: REC-50% and APS-41%, Rgp: REC-61% and APS-67% of the wild type) (Fig. 6A). In contrast, the double null variant (D54A/T271V_FJ_ to D58A/T272V_PG_) had a more pronounced cumulative effect on gingipains secretion resulting in Kgp: 14% and Rgp: 25% of the wildtype (Fig. 6A). Taken together, a clear indication of a cross talk between the REC domain and the APS domain can be seen. Additional analysis of mutants at the dimerization interface revealed that the REC domain dimerization interface (L113E_FJ_ to L117E_PG_) also resulted in reduced gingipain secretion (Kgp: 1% and Rgp: 2% of the wildtype), while the APS domain dimerization interface (S384A/S388E_FJ_ to S385A/S389E_PG_) retaining the gingipains secretion and activity (Kgp: 70% and Rgp: 73% in of the wildtype) (Fig. 6A). These results suggest that maintaining the dimerization ability of the REC domain, and not the APS domain, is of greater significance for the T9SS function. Although both the T272V_PG_ catalytically inactive variant and the S385A/S389E_PG_ APS dimerization interface variant maintained a significant secretion of gingipains, interfering with the Zn1 coordination site residues (D360A/H364A_FJ_ to D361A/H365A_PG_) resulted in reduced gingipain secretion (Kgp: 13% and Rgp: 20% of the wildtype) (Fig. 6A). Based on the ICP-MS results for the D360A/H364A_FJ_ double mutant, which revealed 0.63 zinc ions bound per monomer, it is evident that zinc coordination in Zn1 site has an additional stabilizing effect of on zinc coordination at Zn2 site, presenting with less than a single ion occupancy. Therefore, we conclude that zinc binding at the primary coordination site (Zn1) and associated conformational changes of the 358-369 residues stretch play a more substantial role in the function of the T9SS *in vivo* than the APS domains ability to dimerize.

### PorX_FJ_ mediates digestive enzyme secretion in *F. johnsoniae* but not gliding motility

Deletion of the essential components of T9SS eliminates the crucial function associated with the T9SS, such as, the ability to perform gliding motility and the secretion of digestive enzymes (13, 14, 76). Deletion of *porX* was previously conducted in *F. johnsoniae* strain CJ1827, a streptomycin-resistant *rpsL* mutant of UW101, and it was shown the deletion had no obvious effect on the motility and chitin utilization of *F. johnsoniae* (77). Here, we re-generated the *porX* deletion mutant in the wild-type strain UW101 and examined the phenotypes more quantitively. First, we examined the *ΔporX* effect on the bacterium gliding motility via a classical motility assay and concluded that its genomic deletion does not impact the bacteria’s ability to perform gliding motility (Fig. 6B). Next, we tested the *ΔporX* strain’s ability to grow in a broth media containing chitin as its primary carbon source. Under these restricted conditions, the bacteria must induce and secrete a major chitinase, ChiA, to allow bacterial growth. The dedicated chitinase is secreted by the T9SS and, therefore, serves as a marker for successful digestive enzyme secretion by the T9SS (19). While the deletion of the main T9SS rotary motor and trans-periplasmic complex (*ΔgldKLMNO*) completely abolishes *F. johnsoniae*‘s ability to grow on chitin, the deletion of *porX* results in ∼40% reduced growth and total culture mass (Fig. 6B, lower panel). Overall, the cellular assays demonstrate that PorX does have a mild effect on the T9SS function in *F. johnsoniae*, particularly on the digestive enzyme secretion ability.

## 5. Discussion

Signal transduction by response regulators often requires the phosphorylation and subsequent dimerization of their REC domain (23). Here, we report on the structural determination of PorX, a non-canonical response regulator that comprises a classic REC domain and an additional enzymatic effector domain that exhibits phosphodiesterase activity.

PorX was crystallized in two distinct crystal forms, exhibiting an identical dimeric assembly in both asymmetric units. Therefore, the intertwined dimeric assembly is likely to represent a functional state. Structural analysis and subsequent co-crystallization experiments with phosphate analogs (such as, BeF_3_ or SO4^2-^) established that the observed dimeric assembly represents the phosphorylated-like state and was further confirmed by the positioning of the REC domain’s conserved TY residue pair. PorX’s intertwined dimeric assembly gave rise not only to the classical dimeric interface between two REC domains, but also to an additional interface between two APS domains with both interfaces sharing similar surface area dimensions. The recently determined crystallographic structure of PorX_PG_ revealed a comparable monomeric fold and dimeric assembly to that of PorX_FJ_ (Cα’s RMSDs; monomer = 1.6 Å^2^, dimer 1.9 Å^2^), providing further evidence supporting our conclusion that the intertwined dimer reflects a functional state (Fig. S6) (31).

While the canonical response regulators typically require the phosphorylation of the conserved Asp residue (Asp54 in PorX_FJ_) to induce dimerization of their REC domains, our findings demonstrate that zinc alone can trigger PorX’s dimerization *in vitro*, even in the absence of a potent phosphodonor, like AcP (Fig. 5, Table 1). PorX can bind two zinc cations with high affinity as was demonstrated by biophysical measurements (ICP-MS and ITC). As such, we propose PorX may serve as a zinc sensor, such that altered zinc concentration could constitute one of the dimerization/activation signals for this regulatory cascade and the T9SS. It is important to note that the cellular zinc concentrations are tightly regulated. Studies measuring the total cellular zinc concentration in *E. coli* have demonstrated that the intracellular milieu consists of a myriad of tight zinc-binding proteins (fmol sensitivity), which greatly outnumber the full zinc ion content of the cell (up to 0.2 mM) (78). These findings suggested that under normal growth conditions, there is no persistent pool of free zinc in the cytoplasm (78). Similarly, mammalian cells maintain a total free zinc quota within a narrow range of ∼0.4 fmol/cell, corresponding to a total cellular zinc concentration in the millimolar range (79, 80). While the concentration of free zinc in *P. gingivalis*’s cytosol has yet to be experimentally determined, it is also likely to be low and tightly regulated. Under these conditions, PorX will not constantly exist as a dimer and it is plausible that changes in free zinc concentration trigger, or result from, the activation of the T9SS. Despite the observed nanomolar affinity, we cannot exclude the possibility that the zinc-induced dimerization might be specific to *in vitro* conditions.

**Table 1.**
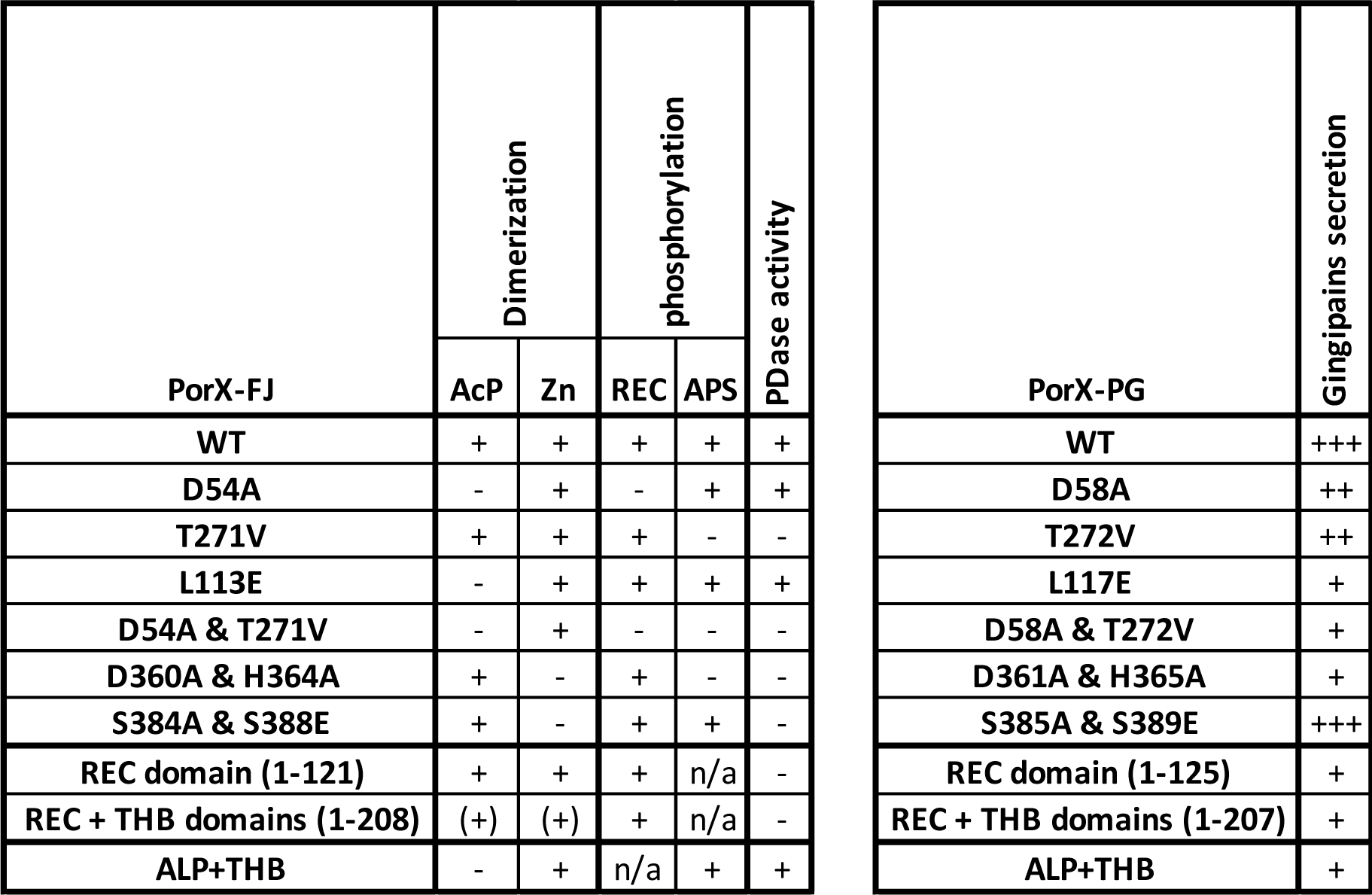
Summary Table. Results of the *in vitro* and *in vivo* characterization of PorX and its variants presented throughout the manuscript. *In vitro* characterization results include oligomeric state analysese by size exclusion chromatography, phosphorylation assays by intact protein MS analysis, and phosphodiesterase (PDase) assays. *In vivo* characterization results include the gingipain activity assays representing the T9SS’s ability to secret gingipains in *P. gingivalis.* The symbols ‘+’, ‘++’ and ‘+++’ in the gingipain secretion column represent average gingipain activities ranging from 0-33%, 34-66%, and 67-100% respectively. The ‘n/a’ indicated that phosphorylation could not be detected as the APS domain was truncated in this construct. The symbol ‘(+)’ indicated that REC+THB domains truncated variant existed as a dimer even before the introduction of dimerizing agent.

PorX’s effector domain adopts the fold of the alkaline phosphatase superfamily. In support of its observed phosphodiesterase activity, the active site coordinates two zinc ions near the catalytic Threonine residue. Structural comparison of the six monomers obtained from our two crystal forms revealed that residues 358-369, near the APS dimerization interface, are of flexible nature as they adopt multiple conformations. Notably, two crucial residues are found within the 358-369 residues stretch, namely Asp360 and His364, that are directly involved in zinc coordination at the APS domain. Therefore, alternation in the conformation of the 358-369 stretch can either establish or abolish the APS’s Zn1 binding site and also affect the stabilization of the bound zinc in Zn2 site (Fig. 1C). Moreover, mutating the Asp360 and His364 dyad was found to affect the APS domain catalytic activity, zinc induced dimerization and the overall T9SS function *in vivo* (Fig. 3, Fig. 5, Table 1).

Although the role of the REC domain in signal transducing by response regulators has been previously established, the role of the APS effector domain *in vivo* remains unclear. Deletion of PorX_PG_’s receiver or effector domains in *P. gingivalis* resulted in a significant decrease in gingipains secretion (<10% of that observed for the wild type), underscoring the contribution of the individual domains to the regulation and function of the T9SS. While the intact APS domain evidently contributes to gingipain secretion, the T272V mutation, which impairs the domain’s catalytic abilities, resulted in only a 50% decrease in gingipain secretion and activity levels, rather than a complete loss (Fig. 6A). Similarly, mutations at the conserved APS dimer interface (S385A/S389E) resulted in a retention of approximately 71% of gingipain secretion (Fig. 6A). In contrast, these APS mutations, when tested *in vitro*, completely abolished phosphodiesterase activity and significantly impaired zinc ion coordination (Fig. 3C, Fig. S5, Table 1). Notably, the D361A/H365A double mutation within the APS domain, which affects the Zn1 binding site coordination and is linked to the conformational changes of residues 358-369, was the only one to cause a significant loss of function both *in vitro* and a reduction in gingipain secretion *in vivo* (Fig. 1C, Table 1). Based on these findings, we conclude that while the enzymatic activity of APS is not essential for signal transduction and the function of the T9SS, the domain’s structural integrity and particularly its zinc binding ability are of significant importance in *P. gingivalis*. Therefore, we propose that the region encompassing residues 358-369 along with its zinc-induced conformational alterations constitute a crucial regulatory site. This site is likely to mediate protein-protein interaction with one or more of the previously reported PorX interacting proteins, such as, SigP, PorL or another protein that must be identified. However, attempts to predict these suggested interactions particularly between PorX dimer, SigP or PorL by AlphaFold (81) did not yield reliable models, as the PorX dimer model always appears to adopt the helical conformation of residues 358-369.

The classical composition of response regulators’ domains often includes an effector domain that directly interacts with DNA. In such cases, the phosphorylation of the REC domain triggers conformational changes, facilitating the effector domain’s binding to DNA, and in particular, to promoter regions. Subsequently, promoter binding induces the transcription of specific gene(s) product(s) and triggers the desired cellular response. Previous biophysical studies report that to compensate for the lack of a DNA-binding effector domain and promote signal transduction throughout the cascade, PorX_PG_ can bind and stabilize the transcription factor, SigP (24). Accordingly, it was shown that only SigP, and not PorX, can bind directly to promoter regions of T9SS-associated genes such as, *porT*, *sov* and *porP* and induce their transcription (24, 26). Other studies have challenged those findings and demonstrated that PorX can also directly bind to promoter regions of T9SS-associted genes, such as *sigP*, *porT*, *PGN_0341* and *PGN_1639* (27). These findings supported the existence of a feedforward loop mechanism involving the direct interaction of both PorX and SigP with the T9SS-associated genes promoters (27). An additional study by the same group further suggested the THB linker domain to serve as the putative DNA-binding region. (28) Here, our findings unambiguously demonstrate that PorX’s effector domain folds and functions as a phosphodiesterase. Moreover, structural comparison and analysis of the entire structure and the THB domain, in particular, revealed no similarity to DNA-binding proteins or transcription factors. Accordingly, PorX’s structure cannot shed light on the reported DNA-binding ability. We speculate that the attributed DNA-binding ability of PorX is mediated by an additional and yet-to-be-identified protein, likely to be involved in this regulatory cascade.

The orphan response regulator PorX was suggested to pair with the orphan SHK, PorY, to form a functional TCS (24, 26). While the genomic deletion of *porX* in *P. gingivalis* abolishes the T9SS function and renders the bacteria avirulent, the genomic deletion of *porY* does not (28, 74). As such, it is plausible that PorX might interact with other SHKs that exist in *P. gingivalis* to render its activity (28, 31, 82). An alternative activation mechanism might also rely on a yet-to-be-identified Asp-kinase or AcP in order to phosphorylate PorX’s REC domain (31), similar to the observed effect of AcP *in vitro* (Fig. 4 & 5). Intriguingly, *in vivo* studies have shown that AcP phosphorylates and activates numerous orphan RR proteins (83). Although the exact concentration of AcP in *P. gingivalis* remains unknown, other bacteria such as *E. coli* have been found to contain at least 3 mM of AcP, which is sufficient for the effective autophosphorylation of RR proteins (84).

PorX is a highly conserved protein found in various bacterial species that utilize their T9SS. Accordingly, our findings demonstrate that PorX_FJ_ can functionally complement the *ΔporX_PG_* strain of *P. gingivalis*. While PorX plays a critical role in the function of the T9SS in the non-gliding *P. gingivalis*, as evidenced by its effects on gingipains secretion and downstream T9SS-associated genes transcription (26–28, 31), its impact on the hallmark T9SS-associated functions in *F. johnsoniae* is different. Here, the deletion of the *porX* gene did not affect the bacteria’s ability to perform gliding motility suggesting that PorX does not regulate the T9SS-attributed gliding motility in *F. johnsoniae*. Next, we assessed the effect of *porX* deletion on the secretion of metabolic enzymes by the T9SS, specifically focusing on the chitinase ChiA. When grown on a chitin-only carbon-source media, the deletion of *porX* gene led to reduced growth compared to the wildtype. However, in both gliding motility and chitinase growth assays, our control deletion of the T9SS-transperilapsmic rotatory motor genes (*gldKLMNO*) completely abolished the gliding motility and the growth on chitin media, respectively. These findings suggest that while PorX is involved in the regulation of T9SS-associated functions in both *P. gingivalis* and *F. johnsoniae*, there are species-specific differences in its impact on the secretion of proteases and polysaccharide degrading enzymes. Further investigation is needed to elucidate the precise mechanisms underlying these differences and the role of PorX in T9SS-related processes in different bacterial species.

Finally, it is important to note that the APS domain of PorX is conserved among T9SS-utilizing bacteria. This raises the possibility that the observed catalytic activity of PorX, in addition to its zinc-induced conformational changes, may have a significant role in the function of PorX in other Bacteroidetes species or in signaling pathways and cellular functions that are not directly related to the T9SS. A recent study has identified a cyclic di-AMP (c-di-AMP) signaling pathway in *P. gingivalis*, which is associated with bacterial growth, regulation of cell envelope homeostasis, and virulence (85). Considering that PorX_PG_ was shown to hydrolyze pApA, a degradation product of c-di-AMP (31), it is plausible that PorX might participate in this signaling pathway. Further investigations are required to explore this possibility and unravel the precise involvement of PorX in other cellular processes and signaling pathways beyond the T9SS.

## 6. Acknowledgments

We thank Dr. Kim Munro from the CRBS for assistance in ITC data collection and the fruitful discussions. We extend our gratitude to Prof. Siavash Vahidi from the University of Guelph, Elaine Wang, Adel Batal and Dr. Simon Veyron from McGill University as well as to Zihe Zhang and Mingbai Zeng from Xi’an Jiaotong-Liverpool University for their assistance during the revision. For mass spectrometry data collection and analysis, the McGill SPR-MS Facility thanks the Canada Foundation for Innovation (CFI) and CRBS for infrastructure support. We thank Dr. Sarah Jantzi at the Plasma Chemistry Laboratory, Center for Applied Isotope Studies, University of Georgia for the ICP-MS data collection. We thank the Canadian Light Source (CLS) beamlines CMCF-ID and CMCF-BM and staff for supporting remote diffraction data collection. The CLS is a national research facility of the University of Saskatchewan, which is supported by the Canada Foundation for Innovation (CFI), the Natural Sciences and Engineering Research Council (NSERC), the National Research Council (NRC), the Canadian Institutes of Health Research (CIHR), the Government of Saskatchewan, and the University of Saskatchewan. We also thank the Advanced Light Source beamline 501 for supporting remote data collection. The Berkeley Center for Structural Biology is supported in part by the National Institutes of Health, the National Institute of General Medical Sciences, and the Howard Hughes Medical Institute. The Advanced Light Source is supported by the Director, Office of Science, Office of Basic Energy Sciences, of the U.S. Department of Energy under Contract No. DE-AC02-05CH11231. This work was funded by the Alzheimer’s Association grant (AARGD-NTF-21-851231) and the Canadian Institutes of Health Research grant to NZ (ARB 1857170). Research reported in this publication was also supported by The National Institute of Dental and Craniofacial Research of the National Institutes of Health under award number R01DE024580 (MED). AS was supported by a CRBS studentship award and the Fonds de Recherche du Québec – Santé (FRQ-S) doctoral fellowship. IM was supported by the Undergraduate Research Center Foundation Grant and Summer Undergraduate Research Experience Grant at Minnesota State University Mankato. JP and MM were supported by Polish National Science Center (UMO-2018/31/B/NZ1/03968 to JP and UMO-2015/19/N/NZ1/00322 to MM). The funders had no role in study design, data collection and analysis, decision to publish, or preparation of the manuscript.

## References

1. G. Hajishengallis, P. I. Diaz, *Porphyromonas gingivalis*: Immune Subversion Activities and Role in Periodontal Dysbiosis. Curr Oral Health Rep 7 (2020).

2. L. M. Sedghi, M. Bacino, Y. L. Kapila, Periodontal Disease: The Good, The Bad, and The Unknown. Front Cell Infect Microbiol 11 (2021).

3. E. Könönen, M. Gursoy, U. K. Gursoy, Periodontitis: A multifaceted disease of tooth-supporting tissues. J Clin Med 8 (2019).

4. M. Kebschull, R. T. Demmer, P. N. Papapanou, “Gum bug, leave my heart alone!”-epidemiologic and mechanistic evidence linking periodontal infections and atherosclerosis. J Dent Res 89, 879–902 (2010).

5. M. Benedyk, et al., Gingipains: Critical Factors in the Development of Aspiration Pneumonia Caused by *Porphyromonas gingivalis*. J Innate Immun 8, 185–198 (2016).

6. O. Laugisch, et al., Citrullination in the periodontium—a possible link between periodontitis and rheumatoid arthritis. Clin Oral Investig 20, 675–683 (2016).

7. S. E. Whitmore, R. J. Lamont, Oral Bacteria and Cancer. PLoS Pathog 10 (2014).

8. S. Gao, et al., Presence of *Porphyromonas gingivalis* in esophagus and its association with the clinicopathological characteristics and survival in patients with esophageal cancer. Infect Agent Cancer 11, 3 (2016).

9. K. Abbayya, N. Y. Puthanakar, S. Naduwinmani, Y. S. Chidambar, Association between periodontitis and Alzheimer’s disease. N Am J Med Sci 7, 241–246 (2015).

10. A. M. Lasica, M. Ksiazek, M. Madej, J. Potempa, The Type IX Secretion System (T9SS): Highlights and Recent Insights into Its Structure and Function. Front Cell Infect Microbiol 7 (2017).

11. N. Li, C. A. Collyer, Gingipains from *Porphyromonas gingivalis* — complex domain structures confer diverse functions. Eur J Microbiol Immunol (Bp) 1 (2011).

12. P. D. Veith, M. D. Glew, D. G. Gorasia, E. C. Reynolds, Type IX secretion: the generation of bacterial cell surface coatings involved in virulence, gliding motility and the degradation of complex biopolymers. Mol Microbiol 106, 35–53 (2017).

13. N. Li, et al., The type IX secretion system is required for virulence of the fish pathogen *Flavobacterium columnare*. Appl Environ Microbiol 83 (2017).

14. P. Barbier, et al., The type IX secretion system is required for virulence of the fish pathogen *Flavobacterium psychrophilum*. Appl Environ Microbiol 86, 1–22 (2020).

15. C. Good, J. Davidson, G. D. Wiens, T. J. Welch, S. Summerfelt, *Flavobacterium branchiophilum* and *F. succinicans* associated with bacterial gill disease in rainbow trout Oncorhynchus mykiss (Walbaum) in water recirculation aquaculture systems. J Fish Dis 38, 409 (2015).

16. Z. Chen, et al., *Riemerella anatipestifer* T9SS Effector SspA Functions in Bacterial Virulence and Defending Natural Host Immunity. Appl Environ Microbiol (2022) 10.1128/AEM.02409-21 (May 23, 2022).

17. B. M. Mark, Y. Zhu, Gliding motility and por secretion system genes are widespread among members of the phylum bacteroidetes. J Bacteriol 195, 270–278 (2013).

18. R. G. Rhodes, S. S. Nelson, S. Pochiraju, M. J. McBride, *Flavobacterium johnsoniae* sprB is part of an operon spanning the additional gliding motility genes sprC, sprD, and sprF. J Bacteriol 193 (2011).

19. S. S. Kharade, M. J. McBride, *Flavobacterium johnsoniae* PorV Is required for secretion of a subset of proteins targeted to the type IX secretion system. J Bacteriol 197 (2015).

20. S. S. Kharade, M. J. McBride, *Flavobacterium johnsoniae* chitinase ChiA is required for chitin utilization and is secreted by the type IX secretion system. J Bacteriol 196 (2014).

21. J. E. Heath, et al., PG1058 is a novel multidomain protein component of the bacterial type IX secretion system. PLoS One 11, 1–29 (2016).

22. F. Jacob-Dubuisson, A. Mechaly, J.-M. M. Betton, R. Antoine, Structural insights into the signalling mechanisms of two-component systems (Nature Publishing Group, 2018).

23. R. Gao, S. Bouillet, A. M. Stock, Structural basis of response regulator function. Annu Rev Microbiol 73, 175–197 (2019).

24. T. Kadowaki, et al., A two-component system regulates gene expression of the type IX secretion component proteins via an ECF sigma factor. Sci Rep 6 (2016).

25. H. Yukitake, et al., PorA, a conserved C-terminal domain-containing protein, impacts the PorXY-SigP signaling of the type IX secretion system. Sci Rep 10, 1–17 (2020).

26. M. S. Vincent, E. Durand, E. Cascales, The PorX response regulator of the *Porphyromonas gingivalis* PorXY two-component system does not directly regulate the type IX secretion genes but binds the PorL subunit. Front Cell Infect Microbiol 6 (2016).

27. C. Jiang, et al., A PorX/PorY and σ P Feedforward Regulatory Loop Controls Gene Expression Essential for *Porphyromonas gingivalis* Virulence. mSphere 6 (2021).

28. D. Yang, C. Jiang, B. Ning, W. Kong, Y. Shi, The PorX/PorY system is a virulence factor of *Porphyromonas gingivalis* and mediates the activation of the type IX secretion system. Journal of Biological Chemistry 296, 1–14 (2021).

29. K. Sato, et al., A protein secretion system linked to bacteroidete gliding motility and pathogenesis. Proceedings of the National Academy of Sciences 107, 276–281 (2010).

30. M. S. Vincent, E. Durand, E. Cascales, The PorX response regulator of the *Porphyromonas gingivalis* PorXY two-component system does not directly regulate the type IX secretion genes but binds the PorL subunit. Front Cell Infect Microbiol 6 (2016).

31. C. Schmitz, et al., Response regulator PorX coordinates oligonucleotide signalling and gene expression to control the secretion of virulence factors. Nucleic Acids Res 50 (2022).

32. S. R. Bond, C. C. Naus, RF-Cloning.org: An online tool for the design of restriction-free cloning projects. Nucleic Acids Res 40 (2012).

33. D. G. Gibson, et al., Enzymatic assembly of DNA molecules up to several hundred kilobases. Nat Methods 6 (2009).

34. F. W. Studier, Protein production by auto-induction in high-density shaking cultures. Protein Expr Purif 41, 207–234 (2005).

35. A. Saran, N. Weerasinghe, C. J. Thibodeaux, N. Zeytuni, Purification, crystallization and crystallographic analysis of the PorX response regulator associated with the type IX secretion system. Acta Crystallogr F Struct Biol Commun 78, 354–362 (2022).

36. Otwinowski Z & Minor W, Processing of X-ray diffraction data collected in oscillation mode. Macromolecular Crystallography, Pt A 276, 307–326 (1997).

37. P. Skubak, et al., A new MR-SAD algorithm for the automatic building of protein models from low-resolution X-ray data and a poor starting model. urn:issn:2052-2525 5, 166–171 (2018).

38. P. Emsley, B. Lohkamp, W. G. Scott, K. Cowtan, Features and development of Coot. Acta Crystallogr D Biol Crystallogr 66, 486–501 (2010).

39. A. J. McCoy, et al., Phaser crystallographic software. urn:issn:0021-8898 40, 658–674 (2007).

40. G. N. Murshudov, et al., REFMAC5 for the refinement of macromolecular crystal structures. Acta Crystallogr D Biol Crystallogr (2011) 10.1107/S0907444911001314.

41. E. F. Pettersen, et al., UCSF Chimera—A Visualization System for Exploratory Research and Analysis. J Comput Chem 25, 1605–1612 (2004).

42. J. F. Trempe, et al., Structural studies of the yeast DNA damage-inducible protein Ddi1 reveal domain architecture of this eukaryotic protein family. Sci Rep 6 (2016).

43. S. Rasool, et al., Mechanism of PINK1 activation by autophosphorylation and insights into assembly on the TOM complex. Mol Cell 82 (2022).

44. M. U. Stevens, et al., Structure-based design and characterization of Parkin activating mutations. bioRxiv (2022).

45. H.-M. Kim, et al., A Novel Regulation of K-antigen Capsule Synthesis in *Porphyromonas gingivalis* Is Driven by the Response Regulator PG0720-Directed Antisense RNA. Frontiers in Oral Health 0, 41 (2021).

46. J. C. Scott, B. A. Klein, A. Duran-Pinedo, L. Hu, M. J. Duncan, A Two-Component System Regulates Hemin Acquisition in *Porphyromonas gingivalis*. PLoS One 8, e73351 (2013).

47. J. Potempa, K. A. Nguyen, Purification and characterization of gingipains. Current protocols in protein science / editorial board, John E. Coligan … [et al.] Chapter 21 (2007).

48. A. M. Lasica, et al., Structural and functional probing of PorZ, an essential bacterial surface component of the type-IX secretion system of human oral-microbiomic *Porphyromonas gingivalis*. Sci Rep 6, 1–22 (2016).

49. M. J. McBride, et al., Novel features of the polysaccharide-digesting gliding bacterium *Flavobacterium johnsoniae* as revealed by genome sequence analysis. Appl Environ Microbiol 75 (2009).

50. M. J. McBride, T. F. Braun, GldI Is a Lipoprotein that Is Required for *Flavobacterium johnsoniae* Gliding Motility and Chitin Utilization. J Bacteriol 186 (2004).

51. M. J. Mcbride, M. J. Kempf, Development of techniques for the genetic manipulation of the gliding bacterium *Cytophaga johnsonae*. J Bacteriol 178, 583–590 (1996).

52. K. D. Cain, B. R. LaFrentz, Laboratory Maintenance of *Flavobacterium psychrophilum* and Flavobacterium columnare. Curr Protoc Microbiol 6 (2007).

53. J. Sambrook, E. F. Fritsch, T. Maniatis, In: Molecular Cloning: A Laboratory Manual, Cold Spring Harbor Laboratory, Cold Spring Harbor, New York. (1989).

54. Y. Zhu, et al., Genetic analyses unravel the crucial role of a horizontally acquired alginate lyase for brown algal biomass degradation by *Zobellia galactanivorans*. Environ Microbiol 19 (2017).

55. D. W. Hunnicutt, M. J. McBride, Cloning and characterization of the *Flavobacterium johnsoniae* gliding motility genes gldD and gldE. J Bacteriol 183 (2001).

56. S. Agarwal, D. W. Hunnicutt, M. J. Mcbride, Cloning and characterization of the *Flavobacterium johnsoniae* (*Cytophaga johnsonae*) gliding motility gene, gldA. Proc Natl Acad Sci U S A 94, 12139–12144 (1997).

57. J. Liu, M. J. McBride, S. Subramaniam, Cell surface filaments of the gliding bacterium *Flavobacterium johnsoniae* revealed by cryo-electron tomography. J Bacteriol 189 (2007).

58. A. Shrivastava, R. G. Rhodes, S. Pochiraju, D. Nakane, M. J. McBride, *Flavobacterium johnsoniae* RemA is a mobile cell surface lectin involved in gliding. J Bacteriol 194 (2012).

59. D. Nakane, K. Sato, H. Wada, M. J. McBride, K. Nakayama, Helical flow of surface protein required for bacterial gliding motility. Proc Natl Acad Sci U S A (2013).

60. M. Dworkin, S. Falkow, E. Rosenberg, K.-H. Schleifer, E. Stackebrandt, The Prokaryotes A Handbook on the Biology of Bacteria (Third Edition Volume 5: Proteobacteria: Alpha and Beta Subclasses) (2006).

61. R. Y. Stanier, The cytophaga group: a contribution to the biology of myxobacteria. Bacteriol Rev 6 (1942).

62. Y. Zhu, et al., Outer membrane proteins related to SusC and SusD are not required for Cytophaga hutchinsonii cellulose utilization. Appl Microbiol Biotechnol 99 (2015).

63. A. Savchenko, et al., Strategies for structural proteomics of prokaryotes: Quantifying the advantages of studying orthologous proteins and of using both NMR and X-ray crystallography approaches. Proteins: Structure, Function, and Bioinformatics 50, 392–399 (2003).

64. R. J. Lewis, J. A. Brannigan, K. Muchová, I. Barák, A. J. Wilkinson, Phosphorylated aspartate in the structure of a response regulator protein. J Mol Biol 294 (1999).

65. V. Agarwal, S. A. Borisova, W. W. Metcalf, W. A. van der Donk, S. K. Nair, Structural and Mechanistic Insights into C-P Bond Hydrolysis by Phosphonoacetate Hydrolase. Chem Biol 18, 1230–1240 (2011).

66. A. Gorelik, F. Liu, K. Illes, B. Nagar, Crystal structure of the human alkaline sphingomyelinase provides insights into substrate recognition. Journal of Biological Chemistry 292, 7087–7094 (2017).

67. J. Morita, et al., Structure and biological function of ENPP6, a choline-specific glycerophosphodiester-phosphodiesterase. Scientific Reports 2016 6:1 6, 1–14 (2016).

68. A. Gorelik, A. Randriamihaja, K. Illes, B. Nagar, A key tyrosine substitution restricts nucleotide hydrolysis by the ectoenzyme NPP5. FEBS J 284, 3718–3726 (2017).

69. R. A. Albright, et al., Molecular Basis of Purinergic Signal Metabolism by Ectonucleotide Pyrophosphatase/Phosphodiesterases 4 and 1 and Implications in Stroke. Journal of Biological Chemistry 289, 3294–3306 (2014).

70. A. Gorelik, A. Randriamihaja, K. Illes, B. Nagar, Structural basis for nucleotide recognition by the ectoenzyme CD203c. FEBS J 285, 2481–2494 (2018).

71. J. Hausmann, et al., Structural basis of substrate discrimination and integrin binding by autotaxin. Nature Structural & Molecular Biology 2011 18:2 18, 198–204 (2011).

72. K. Kato, et al., Crystal structure of Enpp1, an extracellular glycoprotein involved in bone mineralization and insulin signaling. Proc Natl Acad Sci U S A 109, 16876–16881 (2012).

73. J. G. Zalatan, J. G. Zalatan, T. D. Fenn, A. T. Brunger, D. Herschlag, Structural and functional comparisons of nucleotide pyrophosphatase/phosphodiesterase and alkaline phosphatase: Implications for mechanism and evolution. Biochemistry, 9788–9803 (2006).

74. K. Sato, et al., A protein secretion system linked to bacteroidetes gliding motility and pathogenesis. Proceedings of the National Academy of Sciences 107, 276–281 (2010).

75. M. Nonaka, et al., Analysis of a Lys-specific serine endopeptidase secreted via the type IX secretion system in *Porphyromonas gingivalis*. FEMS Microbiol Lett 354 (2014).

76. Y. Zhu, M. J. McBride, Deletion of the *Cytophaga hutchinsonii* type IX secretion system gene sprP results in defects in gliding motility and cellulose utilization. Appl Microbiol Biotechnol 98 (2014).

77. A. Shrivastava, M. J. McBride, Cell surface adhesins, exopolysaccharides and the Por (Type IX) secretion system of *Flavobacterium johnsoniae*. Thesis (2013).

78. C. E. Outten, T. V O’Halloran, Femtomolar sensitivity of metalloregulatory proteins controlling zinc homeostasis. Science 292, 2488–92 (2001).

79. D. A. Suhy, K. D. Simon, D. I. H. Linzer, T. V. O’Halloran, Metallothionein is part of a zinc-scavenging mechanism for cell survival under conditions of extreme zinc deprivation. J Biol Chem 274, 9183–9192 (1999).

80. R. D. Palmiter, S. D. Findley, Cloning and functional characterization of a mammalian zinc transporter that confers resistance to zinc. EMBO J 14, 639–649 (1995).

81. J. Jumper, et al., Highly accurate protein structure prediction with AlphaFold. Nature 596, 583–589 (2021).

82. R. O. Mattos-Graner, M. J. Duncan, Two-component signal transduction systems in oral bacteria. J Oral Microbiol 9 (2017).

83. A. J. Wolfe, Physiologically relevant small phosphodonors link metabolism to signal transduction. Curr Opin Microbiol 13 (2010).

84. A. H. Klein, A. Shulla, S. A. Reimann, D. H. Keating, A. J. Wolfe, The intracellular concentration of acetyl phosphate in Escherichia coli is sufficient for direct phosphorylation of two-component response regulators. J Bacteriol 189 (2007).

85. M. F. Moradali, et al., Atypical cyclic di-AMP signaling is essential for *Porphyromonas gingivalis* growth and regulation of cell envelope homeostasis and virulence. NPJ Biofilms Microbiomes 8 (2022).

