## Supplemental information for "Unveiling the Molecular Mechanisms of the Type-IX Secretion System’s Response Regulator: Structural and Functional Insights"

**Table S1** - Primers used in this study.

| **Primer** | **Sequence (5' --> 3')** | **Purpose** |
|---|---|---|
| *porX*$_{FJ}$ fw | CAGCGGCCTGGTGCCGCGCGGCAGCCATATGGATAAGATAAGAATACTTTGGGTCGATG | pET28a-*porX*$_{FJ}$ |
| *porX*$_{FJ}$ rev | TTGTCGACGGAGCTCGAATTCGGATCCTTATTTAGGGTTAAATACCAAAAACGG | pET28a-*porX*$_{FJ}$ |
| pET28 Kan fw | CCCATTTATACCCATATAAATCAGCATCCATGTTGGAATTTAATCGCGGC | *porX*$_{FJ}$ mutant construction by Gibson assembly |
| pET28 Kan rev | GCCGCGATTAAATTCCAACATGGATGCTGATTTATATGGGTATAAATGGG | *porX*$_{FJ}$ mutant construction by Gibson assembly |
| *porX*$_{FJ}$ L113E fw | ACCAGTAAATCCAAATCAAATTTTAGAGAGTTTAAAAAAGAATCTGGATG | pET28a-*porX*$_{FJ}$ L113E |
| *porX*$_{FJ}$ L113E rev | CATCCAGATTCTTTTTTAAACTCTCTAAAATTTGATTTGGATTTACTGGT | pET28a-*porX*$_{FJ}$ L113E |
| *porX*$_{FJ}$ T271V fw | GTATTTCTCTATTCTTCCAACTGCAGTACAATATGCCAGAAATGCAATTTTC | pET28a-*porX*$_{FJ}$ T271V |
| *porX*$_{FJ}$ T271V rev | GAAAATTGCATTTCTGGCATATTGTACTGCAGTTGGAAGAATAGAGAAATAC | pET28a-*porX*$_{FJ}$ T271V |
| *porX*$_{FJ}$ D54A fw | GAAGAAAACTTTGACATTGTTTTTCTTGCCGAAAATATGCCGGGAATGAGCG | pET28a-*porX*$_{FJ}$ D54A |
| *porX*$_{FJ}$ D54A rev | CGCTCATTCCCGGCATATTTTCGGCAAGAAAAACAATGTCAAAGTTTTCTTC | pET28a-*porX*$_{FJ}$ D54A |
| *porX*$_{FJ}$ K104A fw | CTAAAATCGCAGACTATTTGATAGCACCAGTAAATCCAAATCAAATTTTAC | pET28a-*porX*$_{FJ}$ K104A |
| *porX*$_{FJ}$ K104A rev | GTAAAATTTGATTTGGATTTACTGGTGCTATCAAATAGTCTGCGATTTTAG | pET28a-*porX*$_{FJ}$ K104A |
| *porX*$_{FJ}$ D360A/H364A fw | GTGACTGTTGTTTATAATTTCGTTGCTATGCTTTCGGCTGCAAAAACTGAAATGGAAGTT | pET28a-*porX*$_{FJ}$ D360A/H364A |
| *porX*$_{FJ}$ D360A/H364A rev | AACTTCCATTTCAGTTTTTGCAGCCGAAAGCATAGCAACGAAATTATAAACAACAGTCAC | pET28a-*porX*$_{FJ}$ D360A/H364A |
| *porX*$_{FJ}$ S384A/S388E rev | ATAATGGAGAATTTTTAAACCATTCTAAAGTCAGTGCGCGATATGCTTTGTCATCAGAAG | pET28a-*porX*$_{FJ}$ S384A/S388E |
| *porX*$_{FJ}$ S384A/ S388E fw | CTTCTGATGACAAAGCATATCGCGCACTGACTTTAGAATGGTTTAAAAATTCTCCATTAT | pET28a-*porX*$_{FJ}$ S384A/S388E |

| | | |
|---|---|---|
| *porX*$_{FJ}$ stop codon after 121 fw | ACTGAGTTTAAAAAAGAATCTGGATGATTAATCAAGACTGATTACAGAAAAAAC | pET28a-*porX*$_{FJ}$ REC |
| *porX*$_{FJ}$ stop codon after 121 rev | GTTTTTTCTGTAATCAGTCTTGATTAATCATCCAGATTCTTTTTTAAACTCAGT | pET28a-*porX*$_{FJ}$ REC |
| *porX*$_{FJ}$ stop codon after 208 fw | CTGGTTTGCTCCAAAAGCAGATAAATAACCAATTCAATCTCATAATTTATTTAAAG | pET28a-*porX*$_{FJ}$ REC+THB |
| *porX*$_{FJ}$ stop codon after 208 rev | CTTTAAATAAATTATGAGATTGAATTGGTTATTTATCTGCTTTTGGAGCAAACCAG | pET28a-*porX*$_{FJ}$ REC+THB |
| *porX*$_{FJ}$ 126-end fw | CCGCGCGGCAGCCATATGACAGAAAAAACAACATTAGATTACCAAAAAGAATTC | pET28a-*porX*$_{FJ}$ THB+APS |
| *porX*$_{FJ}$ 126-end rev | CTGGTTTGCTCCAAAAGCAGATAAATAACCAATTCAATCTCATAATTTATTTAAAG | pET28a-*porX*$_{FJ}$ THB+APS |
| FJ0076 | GCTAGGGATCCAACATTATCCCCCAAAACG | Δ*porX*$_{FJ}$ |
| FJ0077 | GCTAGTCTAGAATTGTTGCTTGTTGTAACTTC | Δ*porX*$_{FJ}$ |
| FJ0167 | GCTAGTCTAGATTAGAAGAAATGATTATTCCGTTTT | Δ*porX*$_{FJ}$ |
| FJ0079 | GCTAGGTCGACGATTTTACAGCTGGATAAGAAC | Δ*porX*$_{FJ}$ |
| FJ0086 | GCTAGGGATCCGATGAAAACGTGTATGATTTG | Identification of Δ*porX*$_{FJ}$ CJ4057 |
| FJ0087 | GCTAGTCTAGACAGTCACAGTTTTCACTTCT | Identification of Δ*porX*$_{FJ}$ CJ4057 |
| FJ0088 | GCTAGGGATCCGATGAAAACGTGTATGATTTG | pIM10-*porX* |
| FJ0089 | GCTAGTCTAGACAGTCACAGTTTTCACTTCT | pIM10-*porX* |
| FJ0734 | AATGATGTCGACTCTGCGCTTCCTATTTCG | Δ*gldKLMNO* |
| FJ0735 | CATATCGCATGCTTTCATACGATTTGTATCTGTAGCTGC | Δ*gldKLMNO* |
| FJ1209 | GCTAGGGATCCGCCAATTGCTGTTTACAAAGGAG | Δ*gldKLMNO* |
| FJ1210 | GCTAGGTCGACACCTGACTTACCACAGCCGAT | Δ*gldKLMNO* |
| W50 Δ*porX*$_{PG}$ F1 fw | TTGTAAAACGACGGCCAGTGAATTCTTGGCGACACGCGGACTC | Δ*porX*$_{PG}$::Erm |
| W50 Δ*porX*$_{PG}$ F1 rev | TCTTTTTTGTCATAATTATGTTCTTCTCTATTTAGTATAGGGTATAACGGAAGTG | Δ*porX*$_{PG}$::Erm |
| W50 Δ*porX*$_{PG}$ F2 fw | AAGAACATAATTATGACAAAAAAGAAATTGCC | Δ*porX*$_{PG}$::Erm |

| | | |
|---|---|---|
| W50 Δ*porX*$_{PG}$ F2 rev | TGTATGAAGTATCTACGAAGGATGAAATTTTTC | Δ*porX*$_{PG}$::Erm |
| W50 Δ*porX*$_{PG}$ F3 fw | TCATCCTTCGTAGATACTTCATACATGAATACGATC | Δ*porX*$_{PG}$::Erm |
| W50 Δ*porX*$_{PG}$ F3 rev | CTATGACCATGATTACGCCAAGCTTCAGTATATTGGCC GAATTG | Δ*porX*$_{PG}$::Erm |
| pUC19 sequencing fw | TACGCCAGCTGGCGAAAGGGGGATG | Δ*porX*$_{PG}$::Erm sequencing |
| pUC19 sequencing rev | GCTTTACACTTTATGCTTCCGGCTCG | Δ*porX*$_{PG}$::Erm sequencing |
| W50 Δ*porX*$_{PG}$ fw | ATCACAACGCGAACACCCTGATC | Δ*porX*$_{PG}$::Erm sequencing |
| W50 Δ*porX*$_{PG}$ rev | CCACAGAGGATATATTCGGATAG | Δ*porX*$_{PG}$::Erm sequencing |
| pTMCS groES fw | CGATAAGCTTGGATCCGCATGCCCCATTGGATAGATGC CCTGC | pTCOW-groES-*porX*$_{PG}$ |
| groES PorX$_{PG}$ rev | GTTTTTTTCCATTGTTGCTTGGTTTGTTATTG | pTCOW-groES-*porX*$_{PG}$ |
| groES *porX*$_{PG}$ fw | AAACCAAGCAACAATGGAAAAAAACATGAGACC | pTCOW-groES-*porX*$_{PG}$ |
| *porX*$_{PG}$ pTMCS rev | TAGCGAGGTGCGGCCGGTCGACCCCTTACTTGGGTTGC ATCGTAATTAC | pTCOW-groES-*porX*$_{PG}$ |
| groES *porX*$_{FJ}$ F1 rev | TTATCTTATCCATTGTTGCTTGGTTTGTTATTG | pTCOW-groES-*porX*$_{FJ}$ |
| groES *porX*$_{FJ}$ F2 fw | AACCAAGCAACAATGGATAAGATAAGAATACTTTGG | pTCOW-groES-*porX*$_{FJ}$ |
| *porX*$_{FJ}$ F2 RV | TAGCGAGGTGCGGCCGGTCGACCCCTTATTTAGGGTTA AATACCAAAAAC | pTCOW-groES-*porX*$_{FJ}$ |

| | | |
|---|---|---|
| *porX*$_{PG}$ D58A rev | GTCCGCCGATGCCGGGCATGTTCTCAGCGAGGAATACGATGTCGAAGTCG | pTCOW-groES-*porX*$_{PG}$ D58A |
| *porX*$_{PG}$ D58A fw | CGACTTCGACATCGTATTCCTCGCTGAGAACATGCCCGGCATCGGCGGAC | pTCOW-groES-*porX*$_{PG}$ D58A |
| *porX*$_{PG}$ T272V rev | AGATGGCATTGCGTGCATATTGGACCGCTGTCGGCAGGATGGACAGGTAC | pTCOW-groES-*porX*$_{PG}$ T272V |
| *porX*$_{PG}$ T272V fw | GTACCTGTCCATCCTGCCGACAGCGGTCCAATATGCACGCAATGCCATCT | pTCOW-groES-*porX*$_{PG}$ T272V |
| *porX*$_{PG}$ D361A/H365A fw | GATAGTCCTGAACTTCGTGGCCATGATGTCGGCTGCTCGTACTGATAGCAAGATGATTC | pTCOW-groES-*porX*$_{PG}$ D361A/H365A |
| *porX*$_{PG}$ D361A/H365A rev | GAATCATCTTGCTATCAGTACGAGCAGCCGACATCATGGCCACGAAGTTCAGGACTATC | pTCOW-groES-*porX*$_{PG}$ D361A/H365A |
| *porX*$_{PG}$ S385A/S389E fw | GGCATCCAACGAAGCAGCCTATCGCGCGCTGACGAAGGAATGGTTCAAGCATTCGAC | pTCOW-groES-*porX*$_{PG}$ S385A/S389E |
| *porX*$_{PG}$ S385A/S389E rev | GTCGAATGCTTGAACCATTCCTTCGTCAGCGCGCGATAGGCTGCTTCGTTGGATGCC | pTCOW-groES-*porX*$_{PG}$ S385A/S389E |
| *porX*$_{PG}$ L117E rev | TTTTTTGAGCGACTCGAGGAGCTGATTCGGATTC | pTCOW-groES-*porX*$_{PG}$ L117E |
| *porX*$_{PG}$ L117E fw | TCAGCTCCTCGAGTCGCTCAAAAAAAACCTG | pTCOW-groES-*porX*$_{PG}$ L117E |
| *porX*$_{PG}$ stop codon after 125 | TAGCGAGGTGCGGCCGGTCGACCCCTTACTGCTGCAGGTTTTTTTTG | pTCOW-groES-*porX*$_{PG}$ REC |
| *porX*$_{PG}$ stop codon after 207 | TAGCGAGGTGCGGCCGGTCGACCCCTTACTTGGCAATCCATTCCCGATAG | pTCOW-groES-*porX*$_{PG}$ REC+THB |
| *porX*$_{PG}$ 130-end fw | AACAATGAGCGAAACCACGAACACGAACTACCGGCAAGAGTTCGTCCAAC | pTCOW-groES-*porX*$_{PG}$ THB+APS |
| *porX*$_{PG}$ 130-end rev | CGTGTTCGTGGTTTCGCTCATTGTTGCTTGGTTTGTTATTGTTAGTTGATTGTTTG | pTCOW-groES-*porX*$_{PG}$ THB+APS |
| pTCOW Amp fw | CAGCATCTTTTACTTTCACCAGCGTTTCTGGGTGAGCAAAAACAGGAAGGC | *porX*$_{PG}$ mutant construction by Gibson assembly |

| | | |
|---|---|---|
| pTCOW Amp rev | GCCTTCCTGTTTTTGCTCACCCAGAAACGCTGGTGAAA GTAAAAGATGCTG | *porX*$_{PG}$ mutant construction by Gibson assembly |
| pTCOW seq FW | CAGTGAGGATATTGACGCTTATTTCG | groES-porX sequencing |
| pTCOW seq RV | CGCATTCACAGTTCTCCGCAAG | groES-porX sequencing |

**Table S2** - Strains and plasmids used in this study.

| **Strain (relevant genotype)** | **Source or reference** |
|---|---|
| ***E. coli* strains** | |
| NEB5α | NEB |
| DH5αMCR | Life Technologies |
| BL21 | Life Technologies |
| S17-1 | (1) |
| HB101 | (2, 3) |
| ***F. johnsoniae strains*** | |
| UW101 | (4, 5) |
| UW101 *ΔporX*$_{FJ}$ | This study |
| UW101 *ΔgldKLMNO* | This study |
| ***P. gingivalis* strains** | |
| W50 (wild type) | ATCC |
| *ΔporX*$_{PG}$:Erm (Em$^r$) in strain W50 | This study |
| *ΔporN* in strain W83 | (6) |
| **Plasmids** | |
| pET28a (Kan$^R$) | Novagen |
| pRK2013 (IncP Tra$^+$ Km$^r$) | (3) |
| pYT313 (Ap$^r$ (Em)$^r$ ) | (7) |
| pYT377 (1.9 kb region upstream of *gldK* cloned in pYT313) | This study |
| pYT379 (3 kb region downstream of *gldO* cloned in pYT377) | This study |
| pIM03 (2 kb region upstream of *porX*$_{FJ}$ cloned in pYT313) | This study |
| pIM06 (1.9 kb region downstream of *porX*$_{FJ}$ cloned in pIM03) | This study |
| pCP23 (Ap$^r$ (Tc)$^r$) | (8) |
| pIM10 (2.0 kb region spanning *porX*$_{FJ}$ cloned in pCP23) | This study |
| pUC19 (Amp$^R$) | NEB |
| pT-COW (Amp$^R$ and Tc$^R$ in *E. coli*; Tc$^R$ in *P. gingivalis*; Mob$^+$ Rep$^+$) | (9) |
| pVA2198 (Em$^r$ and Sp$^r$) | (10) |
| pTCOW-groES-*porX*$_{PG}$ | This study |
| pTCOW-groES-*porX*$_{FJ}$ | This study |
| pTCOW-groES-*porX*$_{PG}$ D58A | This study |
| pTCOW-groES-*porX*$_{PG}$ T272V | This study |
| pTCOW-groES-*porX*$_{PG}$ D58A/T272V | This study |
| pTCOW-groES-*porX*$_{PG}$ D361A/H365A | This study |

| | |
|---|---|
| pTCOW-groES-*porX<sub>PG</sub>* S385A/S389E | This study |
| pTCOW-groES-*porX<sub>PG</sub>* L117E | This study |
| pTCOW-groES-*porX<sub>PG</sub>* REC (1-125) | This study |
| pTCOW-groES-*porX<sub>PG</sub>* REC+THB (1-207) | This study |
| pTCOW-groES-*porX<sub>PG</sub>* THB+APS (130-end) | This study |
| pET28a-*porX<sub>FJ</sub>* | This study |
| pET28a-*porX<sub>FJ</sub>* D54A | This study |
| pET28a-*porX<sub>FJ</sub>* T271V | This study |
| pET28a-*porX<sub>FJ</sub>* D54A/T271V | This study |
| pET28a-*porX<sub>FJ</sub>* D360A/H364A | This study |
| pET28a-*porX<sub>FJ</sub>* S384A/S388E | This study |
| pET28a-*porX<sub>FJ</sub>* L113E | This study |
| pET28a-*porX<sub>FJ</sub>* K104A | This study |
| pET28a-*porX<sub>FJ</sub>* REC (1-121) | This study |
| pET28a-*porX<sub>FJ</sub>* REC+THB (1-208) | This study |
| pET28a-*porX<sub>FJ</sub>* THB+APS (126-end) | This study |

**Table S3** - Crystallization conditions and data collection parameters

| | **$PorX_{FJ}$ -$SO_4$** | **$PorX_{FJ}$-Br (also used for phasing)** | **$PorX_{FJ}$-$BeF_3$** | **$PorX_{FJ}$-Zn** | **$PorX_{FJ}$-T271V** |
|---|---|---|---|---|---|
| PDB code | 8TEF | 8TED | 8TFF | 8TFM | 8THP |
| Crystallization condition | 0.2 M Lithium sulfate, 0.1 M Tris pH 8.1 and 35% polyethylene glycol 400 | 0.16 M calcium acetate, 0.08 M sodium cacodylate pH 6.5, 14.4% polyethylene glycol 8000 and 20% glycerol (0.5 M NaBr in the drop) | 0.16 M calcium acetate, 0.08 M sodium cacodylate pH 6.5, 14.4% polyethylene glycol 8000 and 20% glycerol (supplemented with 166 μM beryllium sulphate, 1.162 mM Sodium fluoride and 83 μM Manganese chloride in the drop) | 0.16 M calcium acetate, 0.08 M sodium cacodylate pH 6.5, 14.4% polyethylene glycol 8000, 25% glycerol and 0.2 mM Zinc chloride | 0.16 M calcium acetate, 0.08 M sodium cacodylate pH 6.14, 14.4% polyethylene glycol 8K, 20% Glycerol |

| Beamline | ALS502, Lawrence Berkeley National Laboratory Advanced Light Source (ALS) | CMCF-BM, Canadian Light Source (CLS) | ALS501, Lawrence Berkeley National Laborator y Advanced Light Source (ALS) | CMCF-ID, Canadian Light Source (CLS) | CMCF-ID, Canadian Light Source (CLS) |
|---|---|---|---|---|---|
| Detector | Pilatus3 6M 25 Hz | Pilatus3 6M 25 Hz | Pilatus3 2M 25 Hz | Eiger X 9M | Eiger X 9M |
| Detector distance (mm) | 400 | 347 | 400 | 152.3 | 223.2 |
| Wavelength (Å) | 0.9823 | 0.919 (peak)<br>0.953 (remote) | 0.9774 | 1.283 | 0.953 |
| Oscillation range (°) | 0.25 | 0.5 (peak)<br>0.5 (remote) | 0.25 | 0.2 | 0.2 |
| Time of exposure per image (s) | 0.25 | 2 (peak)<br>1 (remote) | 0.5 | 0.02 | 0.02 |
| Number of images | 1440 | 800 (peak)<br>360 (remote) | 720 | 1800 | 900 |

**Table S4** - Data refinement statistics of the crystals. Statistic values present in parentheses correspond to the highest resolution shell data.

| | **$PorX_{FJ}$-$SO_4$** | **$PorX_{FJ}$-Br (also used for phasing)** | **$PorX_{FJ}$-$BeF_3$** | **$PorX_{FJ}$-Zn** | **$PorX_{FJ}$-T271V** |
|---|---|---|---|---|---|
| PDB code | 8TEF | 8TED | 8TFF | 8TFM | 8THP |
| Space group | $P2_1$ | $P2_12_12_1$ | $P2_12_12_1$ | $P2_12_12_1$ | $P2_12_12_1$ |
| Number of molecules per asymmetric unit | 4 | 2 | 2 | 2 | 2 |
| **Cell dimensions** | | | | | |
| a, b, c (Å) | 133.52, 57.52, 149.14 | 84.70, 97.85, 133.20 | 84.32, 97.52, 131.26 | 86.87, 97.06, 132.04 | 86.09, 97.98, 130.94 |
| α, β, γ (°) | 90.00, 98.39, 90.00 | 90.00, 90.00, 90.00 | 90.00, 90.00, 90.00 | 90.00, 90.00, 90.00 | 90.00, 90.00, 90.00 |
| Resolution (Å) | 46.00-2.85 (2.90-2.85) | 46.16-2.10 (2.15-2.10) | 50.00-2.34 (2.38-2.34) | 50-2.72 (2.77-2.72) | 50-2.6(2.64-2.6) |
| Rsym or Rmerge (%) | 10.30 (32.50) | 8.40 (52.70) | 10.80 (42.20) | 11.3 (82.9) | 9.6 (36.7) |
| R meas (%) | 11.90 (37.90) | 9.90 (62.10) | 11.80 (46.00) | 11.8 (87.0) | 10.8 (40.9) |
| Rpim (%) | 5.80 (19.10) | 5.20 (32.60) | 4.60 (18.20) | 3.3 (25.7) | 4.70(17.9) |
| I / σI | 12.20 (3.85) | 13.70 (3.50) | 17.31 (3.25) | 24.11(2.22) | 18.72 (3.19) |
| Completeness (%) | 100 (100) | 99.60 (99.10) | 100(100) | 99.8(97.7) | 99.9 (99.8) |
| Redundancy | 4 | 6.7 | 6.6 | 12.7 | 5 |
| CC1/2 | 0.984 (0.891) | 0.998 (0.920) | 0.980 (0.910) | 0.998 (0.901) | 0.990 (0.865) |
| CC* | 0.996 (0.971) | 0.999 (0.978) | 0.996 (0.978) | 0.999 (0.974) | 0.998 (0.963) |
| Wavelength (Å) | 0.982 | 0.953 | 0.977 | 1.283 | 0.953 |
| **Refinement** | | | | | |
| Resolution (Å) | 45.40-2.85 | 42.40-2.10 | 48.81-2.34 | 48.58-2.72 | 45.95-2.60 |
| No. reflections (unique) | 53000 | 65014 | 46485 | 31924 | 34201 |
| Rwork / Rfree (%) | 17.67/22.16 | 17.86/22.56 | 16.51/23.10 | 20.43/26.11 | 24.97/30.46 |
| **No. of atoms** | | | | | |
| Protein | 16797 | 8481 | 8476 | 8282 | 8271 |
| Ligand/ion | 59 | 33 | 27 | 18 | 10 |
| Water | 56 | 686 | 381 | 26 | 44 |

| | | | | | |
|---|---|---|---|---|---|
| **B-factors** | | | | | |
| Protein | 51.269 | 33.187 | 38.540 | 43.062 | 52.464 |
| Ligand/ion | 66.654 | 49.76 | 48.698 | 82.374 | 41.502 |
| Water | 27.765 | 37.107 | 35.647 | 50.907 | 25.928 |
| MolProbity score | 1.40 | 1.32 | 1.39 | 1.84 | 1.65 |
| Rotamer outliers (%) | 1.92 | 1.37 | 1.69 | 3.04 | 2.07 |
| Ramachandran favoured (%) | 97.78 | 97.53 | 97.35 | 96.98 | 98.08 |
| Ramachandran allowed (%) | 2.22 | 2.47 | 2.65 | 2.92 | 1.92 |
| Ramachandran outliers | 0 | 0 | 0 | 0.1 | 0 |
| **R.m.s. deviations** | | | | | |
| Bond lengths (Å) | 0.0056 | 0.0084 | 0.0075 | 0.0062 | 0.0057 |
| Bond angles (°) | 1.238 | 1.484 | 1.378 | 1.329 | 1.067 |

**Table S5** – Intact protein LC-MS analyses of $PorX_{FJ}$ variants in the absence or presence of phosphorylation *in vitro*. Expected molecular weights for protein variants (theoretical averages) vs. observed peaks before or after AcP/$Mg^{2+}$ reaction (+ 79.9 single phosphorylation; + 159.8 dual phosphorylation; - 18.0 dehydration; + 61.9 combined dehydration with single phosphorylation).

| | Non-phosphorylated (-AcP) | | Phosphorylated (+AcP) | | | | | | | |
|---|---|---|---|---|---|---|---|---|---|---|
| | | | Single phosphorylation | | Dual phosphorylation | | Cyclization (dehydration) | | Cyclization (dehydration) + single phosphorylation | |
| | Expected | Observed | Expected (+79.9) | Observed | Expected (+159.8) | Observed | Expected (-18.0) | Observed | Expected (-18.0+79.9) | Observed |
| WT | 60449.9 | 60449.1 | 60529.8 | 60528.4 (+79.3) | 60609.7 | 60609.7 (+160.6) | 60431.9 | 60430.9 (-18.2) | 60511.8 | 60511.5 |
| D54A | 60405.9 | 60405.2 | 60485.8 | 60485.5 (+80.3) | – | – | – | – | – | – |
| T271V | 60447.9 | 60447.4 | 60527.8 | 60527.8 (+80.4) | – | – | 60429.9 | 60429.3 (-18.1) | – | – |
| D54A/ T271V | 60403.9 | 60403.5 | – | – | – | – | – | – | – | – |
| D360A/ H364A | 60339.8 | 60338.9 | 60419.7 | 60419.4 (+80.5) | – | – | 60321.8 | 60321.2 (-17.7) | – | – |
| L113E | 60465.8 | 60465.1 60545.4 (at T271)* | 60545.7 | 60545.3 (+80.2) | 60625.6 | 60626.3 (+161.2) | 60447.8 | 60447.2 (-17.9) | 60527.7 | 60527.0 |
| S384A/ S388E | 60475.9 | 60475.9 | 60555.8 | 60555.5 (+79.6) | 60635.7 | 60636.7 (+160.8) | 60457.9 | 60457.7 (-18.2) | 60537.8 | 60537.4 |
| K104A | 60392.8 | 60392.2 | 60472.2 | 60472.8 (+80.6) | – | – | – | – | – | – |
| REC | 14203.4 | 14202.8 | 14283.3 | 14282.7 (+79.9) | – | – | 14185.4 | 14185.1 (-17.7) | – | – |
| REC+THB | 24762.4 | 24761.7 | 24842.3 | 24841.9 (+80.2) | – | – | 24744.4 | 24743.6 (-18.1) | – | – |
| THB+APS | 46208.0 | 46207.0 | 46287.9 | 46286.9 (+79.9) | – | – | – | – | – | – |

*Note: Peak also observed at 60545.4 for L113E variant was likely due to pre-existing phosphorylation at T271 prior to incubation with acetyl phosphate (AcP).

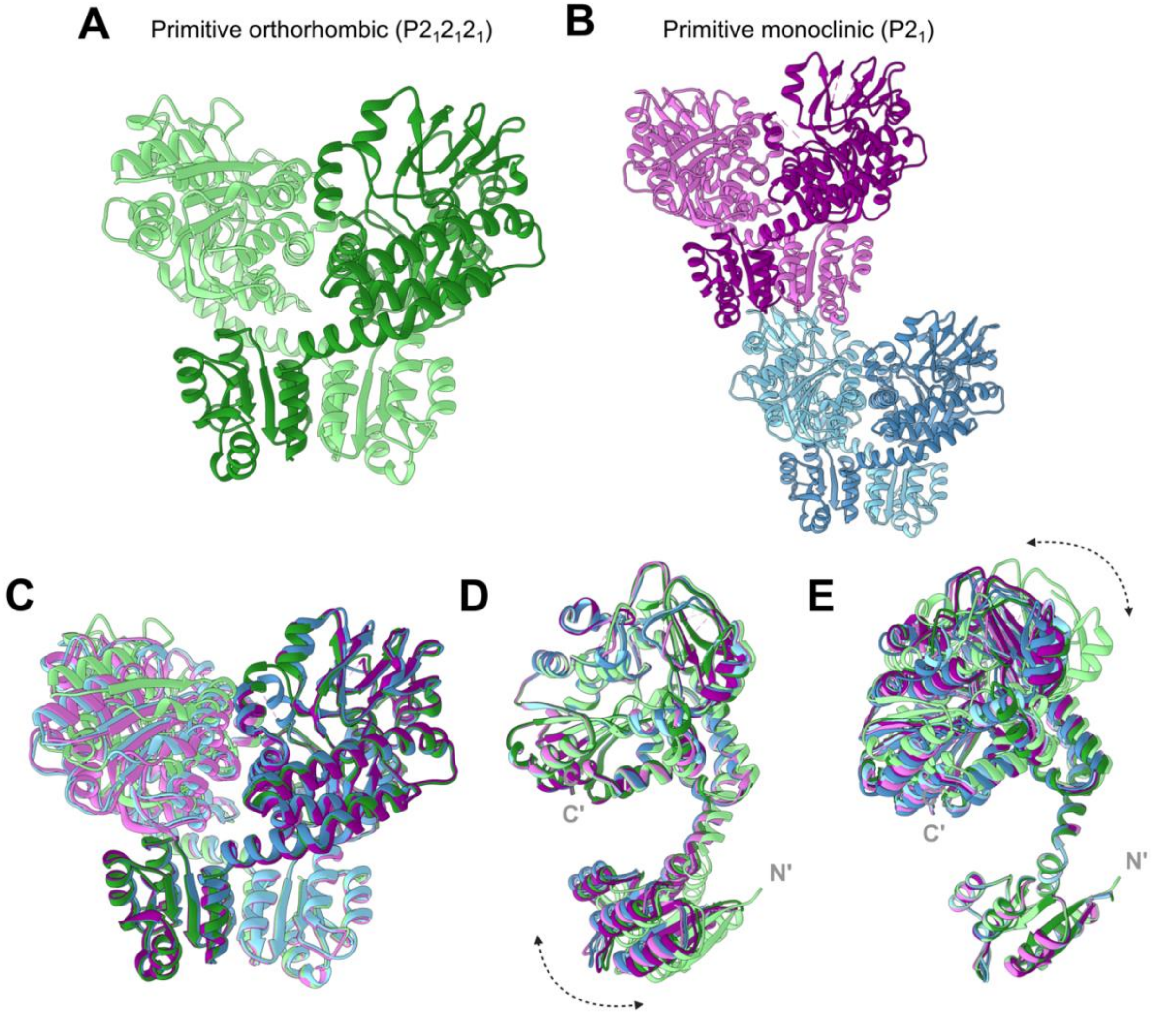


**<u>Fig. S1</u> - PorX$_{FJ}$ structure in two crystal forms.** Asymmetric unit composition of **(A)** primitive orthorhombic and **(B)** primitive monoclinic crystal forms. **(C)** Superposition of the three dimers shown in panels A and B. Overlay of the six monomers shown in A and B aligned according to their **(D)** alkaline phosphatase superfamily (APS) domain and **(E)** receiver (REC) domain.

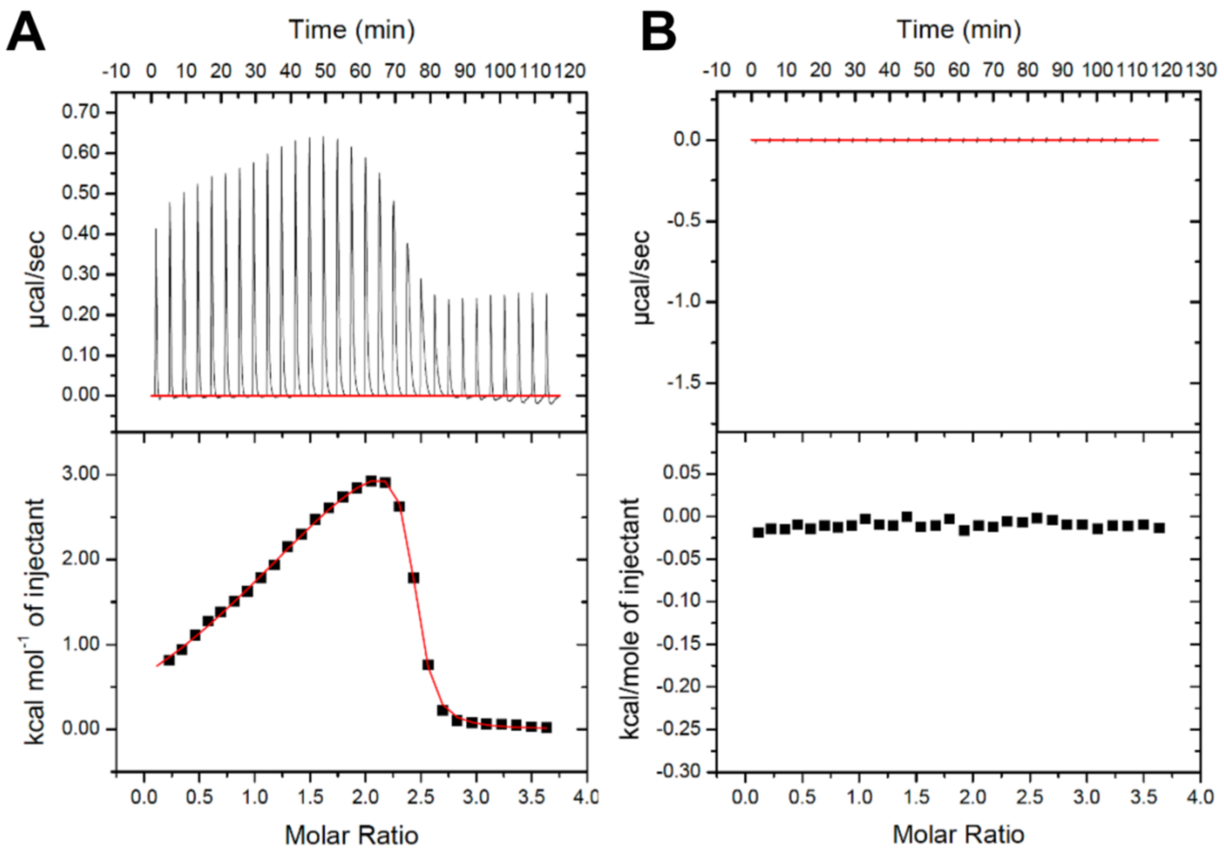


**Fig. S2** - **Zinc binding to PorX$_{FJ}$ by isothermal titration calorimetry. (A)** Binding isotherm of zinc titrated into PorX$_{FJ}$ reveals two zinc sites stoichometry with $N_1$= 1.09 ± 0.04 sites, $K_{D1}$=180.83 ± 14.93 nM, $\Delta H_1$=-2.374E4 ± 1.28E6 cal/mol, $\Delta S_1$=-50.2 cal/mol/deg and $N_2$=1.19 ± 0.04 sites, $K_{D2}$= 59.17 ± 13.81 nM, $\Delta H_2$=2.744E4 ± 1.26E6 cal/mol, $\Delta S_2$=124 cal/mol/deg. **(B)** Buffer titrated into PorX$_{FJ}$ as a control.

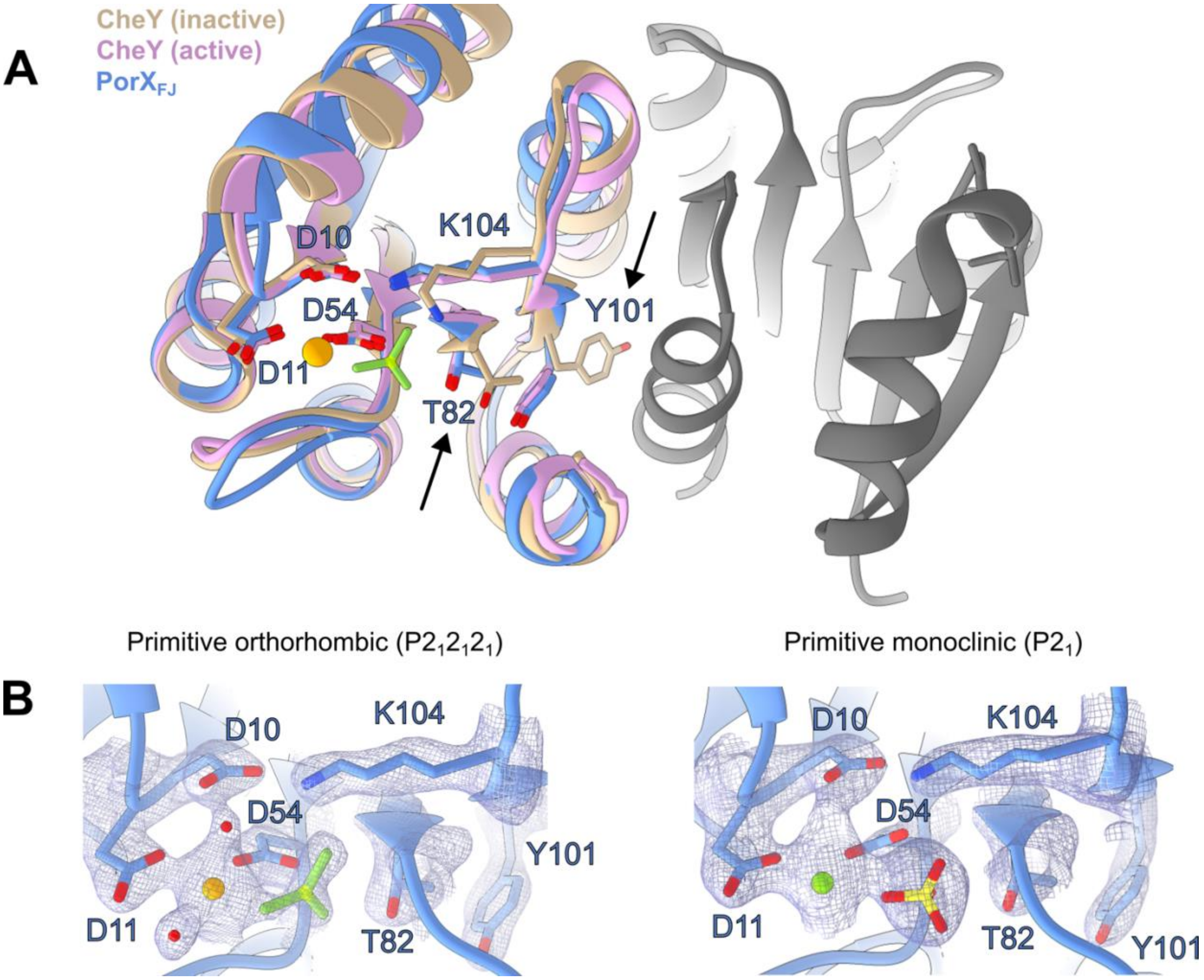


**<u>Fig. S3</u>** – The crystal structure of PorX$_{FJ}$ adopts the active phosphorylated-like conformation. **(A)** Superposition of the primitive orthorhombic crystal form of PorX$_{FJ}$ with the active (PDB ID:1FQW) and inactive (PDB ID: 2CHE) conformations of the CheY response regulator reveal an active-like conformation of PorX$_{FJ}$, as per the positioning of the conserved Thr-Tyr pair, marked by arrows. Inactive and active conformations of CheY are colored in tan and pink, while chains A and B of PorX$_{FJ}$ dimer are colored in blue and grey, respectively. Specific residue labels are according to PorX$_{FJ}$ sequence. **(B)** Electron density of phosphate analogs, $BeF_3$ and $SO_4^{2-}$, at the REC domain phosphorylation site in the different crystal forms of PorX$_{FJ}$. Orange, green and red spheres represent $Ca^{2+}$, $Mg^{2+}$ and $H_2O$ respectively.

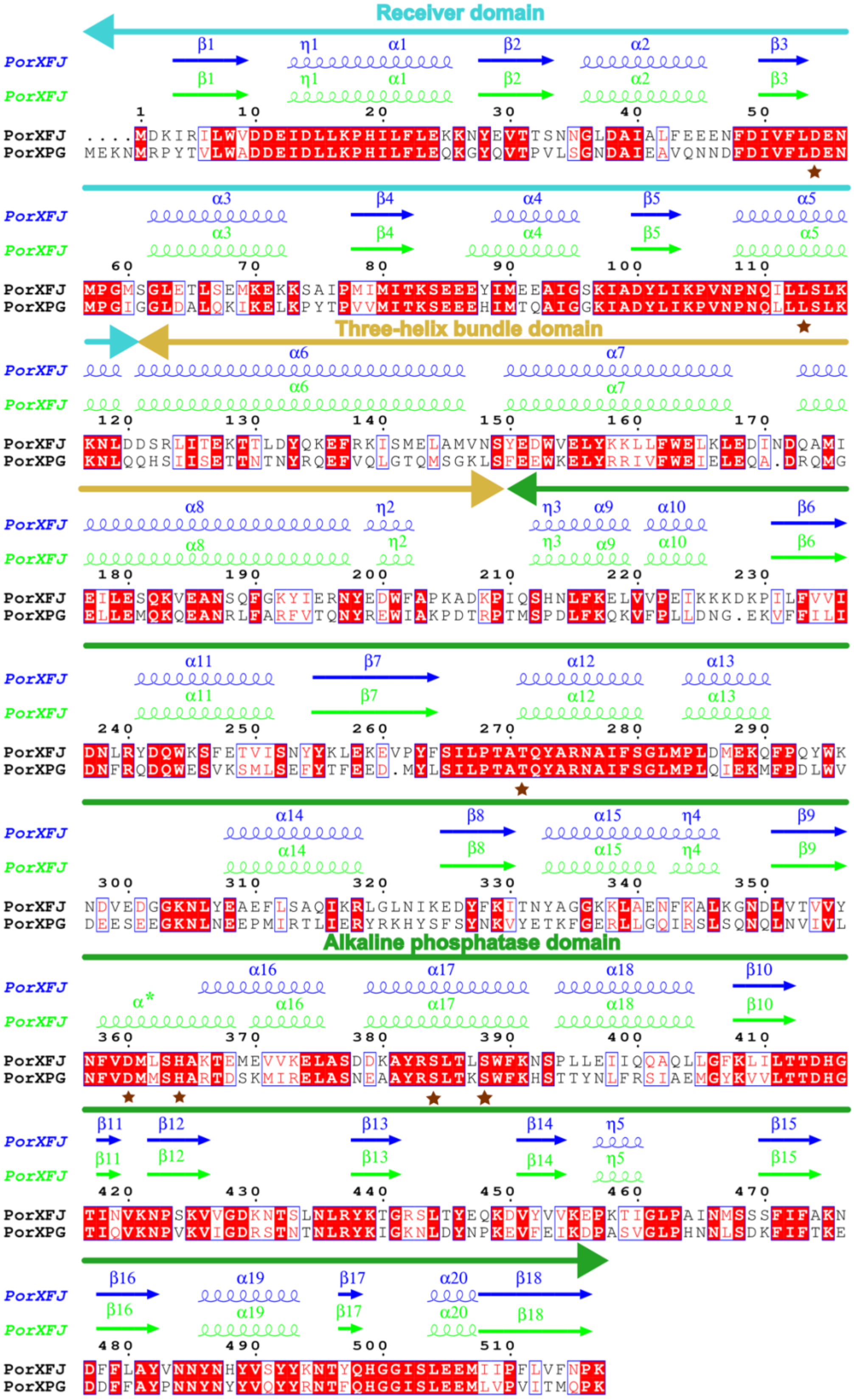

Receiver domain
Three-helix bundle domain
Alkaline phosphatase domain
PorXFJ
PorXPG

**Fig. S4** - **Multiple sequence alignment of PorX$_{FJ}$ and PorX$_{PG}$**. The alignment was prepared using Clustal Omega (11) and ESPript (12). The receiver, three-helix bundle and alkaline phosphatase domains are labelled in cyan, yellow and green respectively. Reference amino acid numbering and secondary structure prediction is according to the PorX$_{FJ}$ sequence. The secondary structure alignment in blue and light green correspond to the alternate chain conformations of the primitive orthorhombic dimeric structure with α* indicating the area of conformational difference between the two chains. Amino acid residues marked with a star correspond to functionally significant residues whose mutations are discussed in this manuscript.

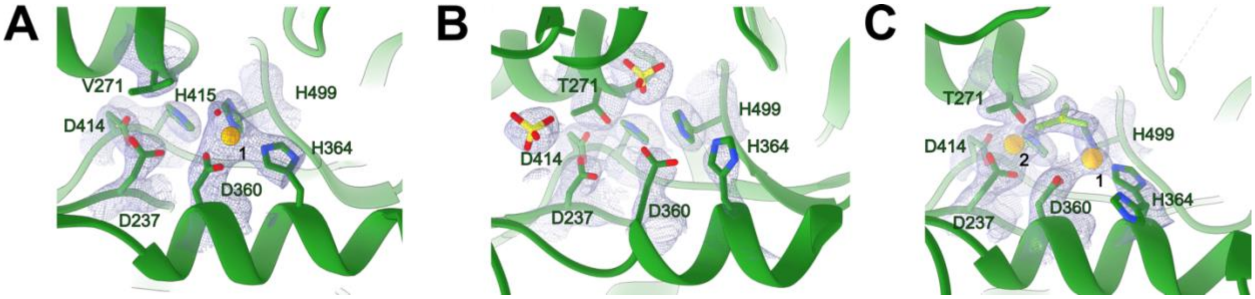


**<u>Fig. S5</u>** – Electron density of divalent cations and phosphate analogs in the APS domain of **(A)** PorX$_{FJ}$-T271V crystal form **(B)** PorX$_{FJ}$-$SO_4^{2-}$ primitive monoclinic crystal form and **(C)** PorX$_{FJ}$-$BeF_3$ primitive orthorhombic crystal form**.**

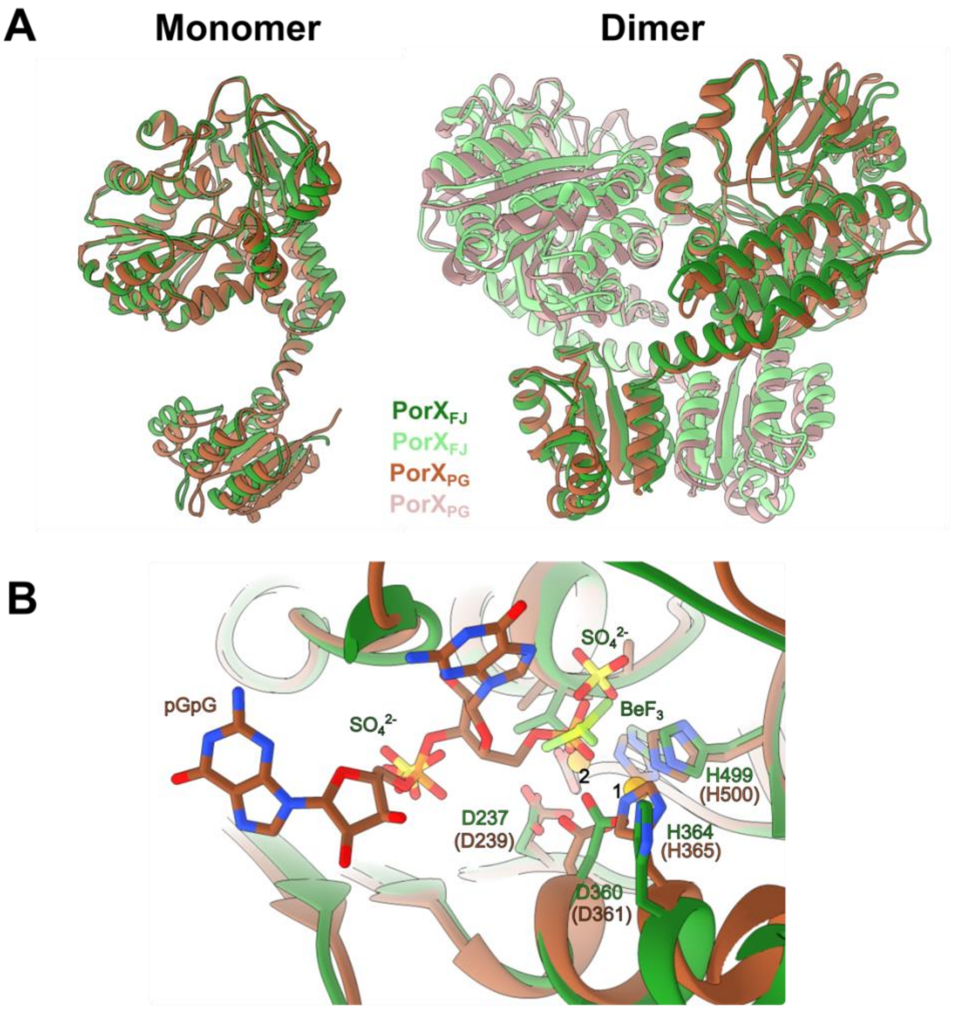


**Fig. S6 – The structures of PorX$_{FJ}$ and PorX$_{PG}$ (PDB code:7PVK) exhibit a highly conserved fold. (A)** A superimposition of a representative monomer and a representative dimeric assembly. **(B)** Superimposition of the active site of the APS domain. Ligand analogs $BeF_3$ and sulphate ions observed in the PorX$_{FJ}$ structures directly overlap or are situated in close proximity to the phosphoguanylyl-(3′→5′)-guanosine (pGpG) bound to PorX$_{PG}$.

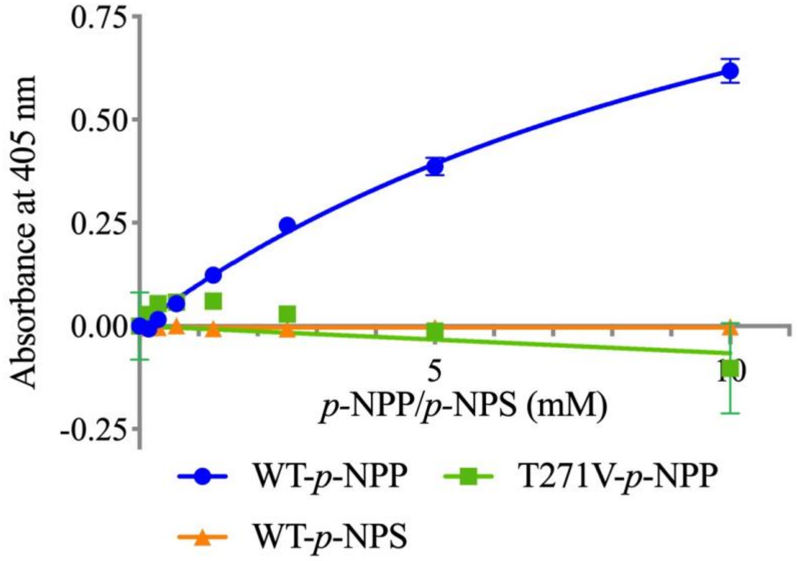


**Fig. S7** - **Monophosphatase and sulphatase activities of PorX$_{FJ}$.** Catalytic activities against *p*-nitrophenyl phosphate (*p*-NPP) and *p*-nitrophenyl sulphate (*p*-NPS) substates were recorded in the presence of zinc after three days.

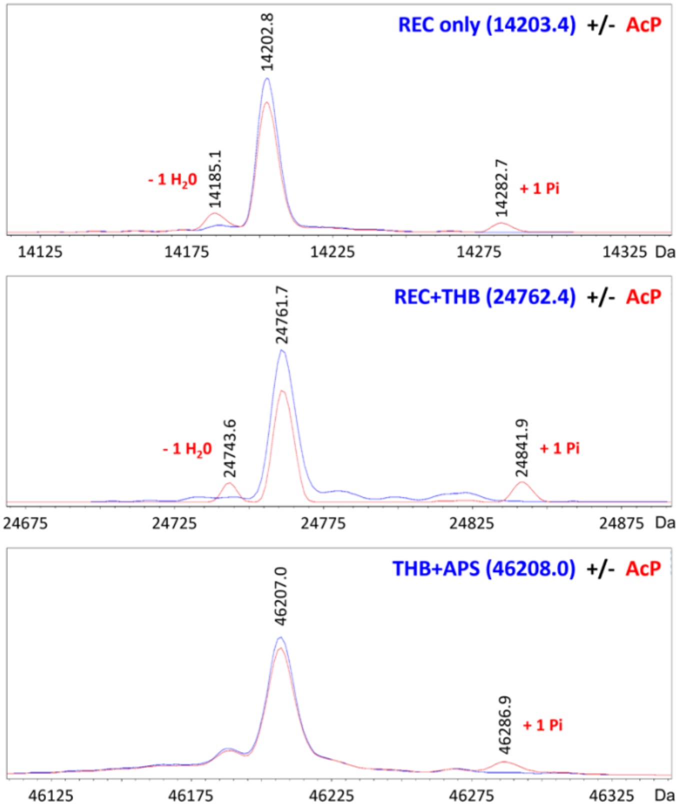


**<u>Fig. S8</u> - Intact protein LC-MS analyses of PorX$_{FJ}$ truncation variants in the absence or presence of phosphorylation *in vitro***. The blue and red spectra correspond to non-phosphorylated and phosphorylated versions of the protein, respectively. The acetyl phosphate (AcP) phosphodonor promotes the phosphorylation of Asp54 and/or Thr271 ("+1 Pi" label). Alternatively, AcP was found to induce cyclization of Asp54 and Lys104, resulting in a dehydration reaction ("-1 $H_2$0" labels).

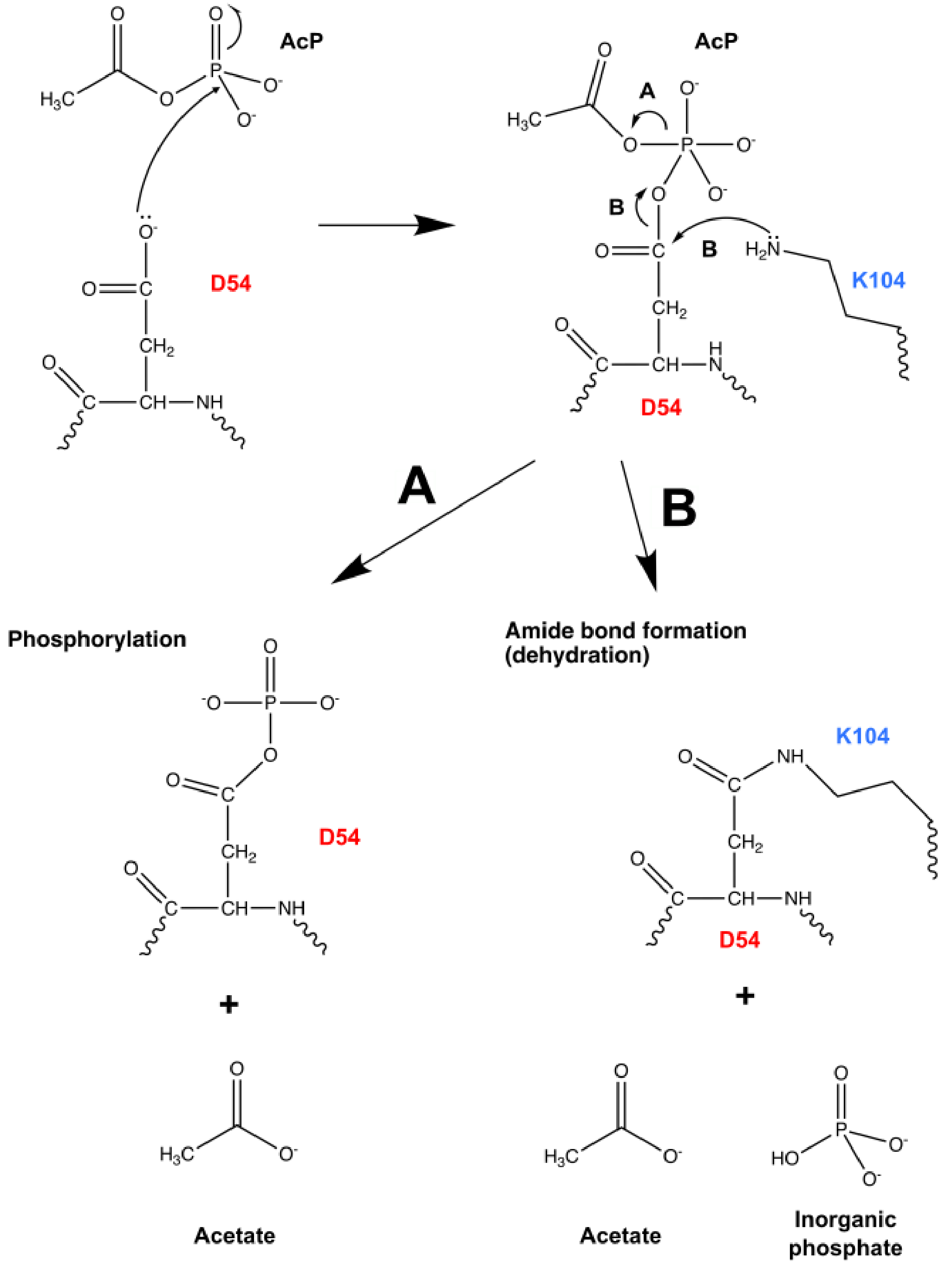


**<u>Fig. S9</u> - Proposed phosphorylation mechanism by acetyl phosphate *in vitro*.** Reaction of active D54 with AcP can lead to **(A)** phosphorylation and/or **(B)** cyclization involving the formation of an internal peptide linkage between D54 and K104.

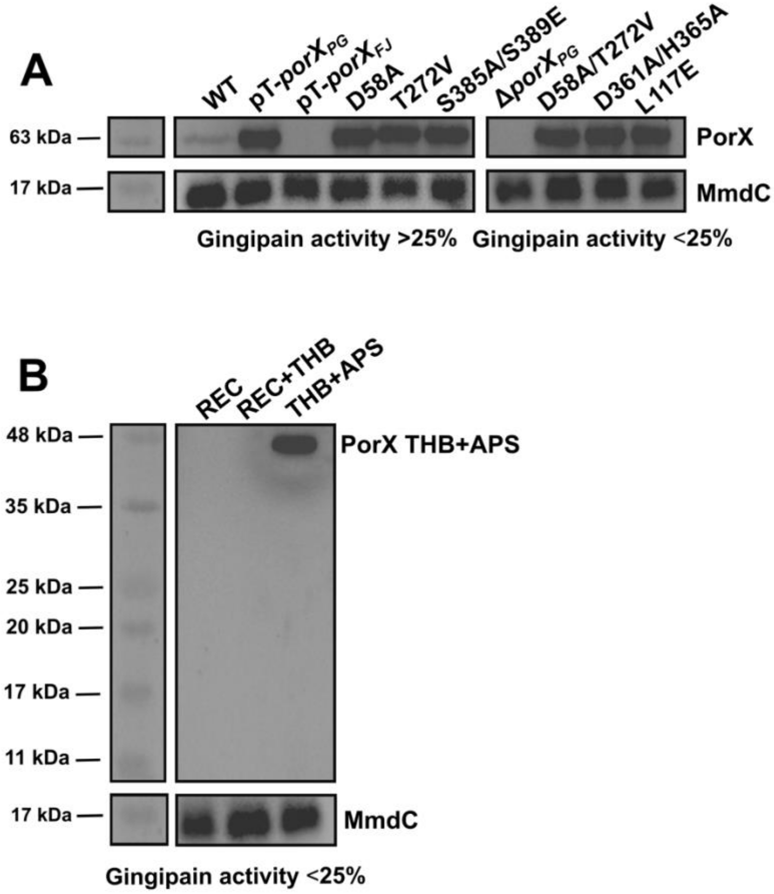


**<u>Fig. S10</u> - Expression levels of PorX in *P. gingivalis***. (A) point mutations and (B) truncation variants. Whole cell lysates underwent SDS-PAGE separation followed by Western blot analysis. A rabbit polyclonal anti-PorXPG primary antibody and a polyclonal goat anti-rabbit horseradish peroxidase-conjugated secondary antibody were employed to detect PorX (~61 kDa). The biotinylated protein MmdC (~15 kDa) was used as a loading control (13) and was detected using horseradish peroxidase conjugated Streptavidin.

The anti-PorX$_{PG}$ did not recognize PorX$_{FJ}$ due to sequence variations that affect epitope recognition. Among the truncation variants, PorXPG-REC (~14 kDa) and PorX$_{PG}$-REC+THB (~24 kDa) were not detected, while the PorX$_{PG}$-THB+APS variant (~46 kDa) was identified by the anti-PorX$_{PG}$. This selective recognition of the APS domain truncation variant by the anti-PorX$_{PG}$ antibody may be attributed to the presence of recognizable epitopes exclusively within the APS domain. Nonetheless, the possibility of low expression levels or the instability and subsequent degradation of the PorX$_{PG}$-REC and PorX$_{PG}$-REC+THB truncation variants cannot be excluded.

**References:**


1. C. Matsumoto-Mashimo, A.-M. Guerout, D. Mazel, A new family of conditional replicating plasmids and their cognate Escherichia coli host strains. *Research in microbiology* **155**, 455-461 (2004).
2. F. Bolivar, K. Backman, "[16] Plasmids of Escherichia coli as cloning vectors" in Methods in enzymology. (Elsevier, 1979), vol. 68, pp. 245-267.
3. D. H. Figurski, D. R. Helinski, Replication of an origin-containing derivative of plasmid RK2 dependent on a plasmid function provided in trans. *Proceedings of the National Academy of Sciences* **76**, 1648-1652 (1979).
4. M. J. McBride *et al.*, Novel features of the polysaccharide-digesting gliding bacterium Flavobacterium johnsoniae as revealed by genome sequence analysis. *Applied and environmental microbiology* **75**, 6864-6875 (2009).
5. M. J. McBride, T. F. Braun, GldI is a lipoprotein that is required for Flavobacterium johnsoniae gliding motility and chitin utilization. *Journal of bacteriology* **186**, 2295-2302 (2004).
6. A. M. Lasica *et al.*, Structural and functional probing of PorZ, an essential bacterial surface component of the type-IX secretion system of human oral-microbiomic Porphyromonas gingivalis. *Sci Rep* **6**, 37708 (2016).
7. Y. Zhu *et al.*, Genetic analyses unravel the crucial role of a horizontally acquired alginate lyase for brown algal biomass degradation by Z obellia galactanivorans. *Environmental microbiology* **19**, 2164-2181 (2017).
8. S. Agarwal, D. W. Hunnicutt, M. J. McBride, Cloning and characterization of the Flavobacterium johnsoniae (Cytophaga johnsonae) gliding motility gene, gldA. *Proceedings of the National Academy of Sciences* **94**, 12139-12144 (1997).
9. R. G. Gardner, J. B. Russell, D. B. Wilson, G.-R. Wang, N. B. Shoemaker, Use of a modified Bacteroides-Prevotella shuttle vector to transfer a reconstructed beta-1, 4-D-endoglucanase gene into Bacteroides uniformis and Prevotella ruminicola B (1) 4. *Applied and environmental microbiology* **62**, 196-202 (1996).
10. H. M. Fletcher *et al.*, Virulence of a Porphyromonas gingivalis W83 mutant defective in the prtH gene. *Infection and immunity* **63**, 1521-1528 (1995).
11. F. Madeira *et al.*, Search and sequence analysis tools services from EMBL-EBI in 2022. *Nucleic Acids Research* **50**, W276-W279 (2022).
12. X. Robert, P. Gouet, Deciphering key features in protein structures with the new ENDscript server. *Nucleic Acids Research* **42**, W320-W324 (2014).
13. A. M. Lasica *et al.*, Structural and functional probing of PorZ, an essential bacterial surface component of the type-IX secretion system of human oral-microbiomic Porphyromonas gingivalis. *Scientific Reports* **6**, 37708 (2016).
